# Slow-rising and fast-falling dopaminergic dynamics jointly adjust negative prediction error in the ventral striatum

**DOI:** 10.1101/2021.07.23.453499

**Authors:** Yu Shikano, Sho Yagishita, Kenji F. Tanaka, Norio Takata

## Abstract

The greater the reward expectations are, the more different the brain’s physiological response will be. Although it is well-documented that better-than-expected outcomes are encoded quantitatively via midbrain dopaminergic (DA) activity, it has been less addressed experimentally whether worse-than-expected outcomes are expressed quantitatively as well. We show that larger reward expectations upon unexpected reward omissions are associated with the preceding slower rise and following larger decrease (DA dip) in the DA concentration at the ventral striatum of mice. We set up a lever press task on a fixed ratio (FR) schedule requiring five lever presses as an effort for a food reward (FR5). The mice occasionally checked the food magazine without a reward before completing the task. The percentage of this premature magazine entry (PME) increased as the number of lever presses approached five, showing rising expectations with increasing proximity to task completion, and hence greater reward expectations. Fiber photometry of extracellular DA dynamics in the ventral striatum using a fluorescent protein (genetically encoded GPCR-activation-based-DA sensor: GRAB_DA2m_) revealed that the slow increase and fast decrease in DA levels around PMEs were correlated with the PME percentage, demonstrating a monotonic relationship between the DA dip amplitude and degree of expectations. Computational modeling of the lever press task implementing temporal difference errors and state transitions replicated the observed correlation between the PME frequency and DA dip amplitude in the FR5 task. Taken together, these findings indicate that the DA dip amplitude represents the degree of reward expectations monotonically, which may guide behavioral adjustment.

## Introduction

Learning an appropriate association between behavior and outcome is critical for animal survival. Reinforcement learning (RL) algorithms are proposed as neuronal implementations for such experience-based learning (Schultz et al., 1997). Such an algorithm updates the subjective value of a state or an action based on the difference between the expected reward and the actual reward obtained, which is called the reward prediction error (RPE). An increasing number of studies have demonstrated that a better-than-expected outcome (i.e., a positive RPE) is encoded quantitatively by spiking activity of midbrain dopaminergic (DA) neurons (Bayer and Glimcher, 2005; Tobler et al., 2005) and by extracellular DA levels in the ventral striatum (Covey and Cheer, 2019; Heien et al., 2005). In contrast, there are still ongoing debates regarding the neuronal representation of the magnitude of a worse-than-expected outcome inducing a negative RPE **(*Hart et al., 2014*)**. While positive and negative RPEs are treated in the same way in RL algorithms, it is not clear whether the neuronal implementations of negative RPEs are similar to those of positive RPEs. Indeed, the economic expected utility theory (Neumann and Morgenstern, 1945) and the psychological prospect theory (Kahneman and Tversky, 1979) claim that the subjective perceptions of positive and negative rewards are *a*symmetric. Additionally, a biological investigation revealed that a DA surge and decrease in positive and negative RPEs, respectively, evoke distinct biochemical responses in different downstream targets (Iino et al., 2020). Furthermore, it is not clear when the DA level modulation occurs to realize the monotonic variations in the DA dip amplitude for reflecting the magnitude of negative RPEs upon unexpected reward omissions. While longer inter-spike intervals of DA neurons might realize deeper trough levels of DA fluctuations (Bayer et al., 2007), it could be a pre-dip DA level rather than the dip-trough level that constitutes variations of the DA dip amplitude (Hamid et al., 2016).

DA dynamics in the ventral striatum have been investigated using an electrophysiological measurement of midbrain DA neurons that send afferents to the ventral striatum. Whereas firing activity of the DA neurons can be observed with a high temporal resolution, DA release dynamics in the ventral striatum cannot be determined, as spiking-activity-independent DA release has been identified (Mohebi et al., 2019). Fast scan cyclic voltammetry (FSCV) has been a standard method for the direct detection of extracellular DA, extending our knowledge of DA dynamics during behavior (Heien et al., 2005; Phillips et al., 2003; Robinson et al., 2003; Rodeberg et al., 2017). Recently, a genetically encoded optical DA sensor, G protein-coupled receptor-activation-based DA (GRAB_DA2m_) was reported (Sun et al., 2018; Sun et al., 2020). GRAB_DA2m_ has a high sensitivity (EC_50_ values of 90 nM), high specificity (10‒20-fold more selective for DA over structurally similar noradrenaline) and long-term stability (> 120 min), suited for addressing the debate on a DAergic representation of a negative RPE (Hamid et al., 2016).

In the present study, we examined the neuronal representation of reward expectations using a DA signal in the ventral striatum. We developed a fiber photometry system equipped with a highly sensitive Gallium Arsenide Phosphide (GaAsP) photocathode and combined this system with the viral expression of GRAB_DA2m_ proteins in the ventral striatum of mice. We monitored the DA dynamics during a lever press task on a fixed ratio (FR) schedule. The mice had to press a lever five times to obtain a food reward (FR5) (Yoshida et al., 2019). The advantage of this simple operant task is that it brought about various degrees of reward expectations to the mice 1) using a fixed reward size (always a single pellet) after the completion of a FR5 task, and 2) during almost identical behavior patterns in anticipation of a reward. These characteristics contrast with other popular tasks that use variable reward sizes (e.g., the amount of juice) or forces a behavioral choice (e.g., left or right) for modulating the magnitude of expectation—either of which may introduce extra DA dynamics not related to reward expectations. We found that the larger reward expectations, which was estimated based on behavior, are associated with the preceding higher baseline DA level and the following deeper DA dip in the ventral striatum. An RL model using the temporal-difference (TD) algorithm (Amo et al., 2020; Schultz et al., 1997) supported that the modulation of the DA dip amplitude according to the degree of reward expectations was important for the mice to establish adaptive behavior in the FR5 task. Taken together, we propose that the degree of reward expectations is represented quantitatively by the magnitude of the DA dip, which plays a role in behavior adjustment.

## Materials and Methods

### Animals

All animal procedures were conducted in accordance with the National Institutes of Health Guide for the Care and Use of Laboratory Animals and approved by the Animal Research Committee of Keio University School of Medicine (approval 14027-(1)). Experiments were performed using three-month-old male C57BL/6JJmsSlc or C57BL/6NCrSlc mice weighing 23–27 g purchased from SLC (Shizuoka, Japan). The mice were housed individually and maintained on a 12-h light/12-h dark schedule, with lights off at 8:00 PM. Their body weights were maintained at 85% of their initial body weight under conditions of food restriction with water ad libitum.

### Adeno-Associated Virus Preparation

For adeno-associated virus (AAV) production, we used the pAAV-hSyn-GRAB-DA2m-W plasmid, a gift from Y. Li. We produced the AAV with the PHP.eB serotype, allowing for the efficient transduction of the central nervous system (Mathiesen et al., 2020). Its titer was measured as described elsewhere (Grieger et al., 2006). In brief, the plasmid, pHelper (Stratagene), and PHP.eB (Addgene, Plasmid #103005) were transfected into HEK293 cells (AAV293, Stratagene). After three days, the cells were harvested, and the AAV was purified twice using iodixanol. The titer for the AAV was estimated via quantitative PCR.

### Surgery

A total of eight mice underwent stereotaxic surgery under anesthesia with a ketamine-xylazine mixture (100 and 10 mg/kg, respectively, i.p.). A 8-mm midline incision was made from the area between the eyes to the cerebellum. A craniotomy with a diameter of up to 1.5 mm was created above the right ventral striatum (VS) (1.1 mm anterior and 1.9 mm lateral to the bregma) using a high-speed drill, and the dura was surgically removed. A total volume of 0.5 μL GRAB_DA2m_ virus (PHP.eB AAV-hSyn-GRAB-DA2m-W, 1.0 × 10^14^ genome copies/ml) was injected through a pulled glass micropipette into the VS (−3.5 to −3.7 mm ventral from dura surface) according to the atlas of Paxinos and Franklin (2004). The injection was driven at a flow rate of 100 nL/min by a Nanoliter 2020 Injector (World Precision Instruments, Sarasota, FL). The micropipette was left in place for another 5 minutes to allow for tissue diffusion before being retracted slowly. Following the GRAB_DA2m_ virus injection, an optical fiber cannula (CFMC14L05, 400 mm in diameter, 0.39 NA; Thorlabs, Newton, NJ) attached to a ceramic ferrule (CF440-10, Thorlabs) and a ferrule mating sleeve (ADAF1-5, Thorlabs) were inserted into the same side of the VS as the virus injection and cemented in place (−3.4 to −3.6 mm ventral from dura surface). Operant conditioning and data collection were started in more than 10 days after the surgery to allow the mice to recover and the GRAB_DA2m_ proteins to be expressed.

To confirm that neither GRAB_DA2m_ expression nor the optic fiber implantation altered mice’s behavior in our behavior task, another group of six mice underwent stereotaxic surgery under anesthesia, and a midline incision was made as described in the paragraph above. For four of these mice, no craniotomy was made, and the skull was cemented. The other two mice received saline injections bilaterally (0.2 μL/site) targeting the VS through the craniotomies. The skull was cemented.

### Fiber Photometry and Signal Extraction

Extracellular DA fluctuations were measured by a fiber photometric system designed by Olympus Corporation, Tokyo, Japan. Extracellular DA fluorescence signals were obtained by illuminating cells expressing GRAB_DA2m_ with a 465 nm LED (8.0 ± 0.1 μW at the fiber tip) and a 405 nm LED (the same power as that of the 465 nm LED). The 405 nm LED was used to correct for movement artifacts. The 465 nm and 405 nm LED light were emitted alternately at 20 Hz (turned on for 24 ms and off for 26 ms), with the timing precisely controlled by a programmable pulse generator (Master-8, A.M.P.I., Jerusalem, ISRAEL). Each excitation light was reflected by a dichroic mirror (DM455CFP; Olympus) and coupled into an optical fiber cable (400 μm in diameter, 2 meters in length, 0.39 NA, M79L01; Thorlabs, Newton, NJ) through a pinhole (400 μm in diameter). The optical fiber cable was connected to the optical fiber cannula of the mice. The fluorescence signal was detected by a photomultiplier tube with a GaAsP photocathode (H10722–210; Hamamatsu Photonics, Shizuoka, Japan) at a wavelength of 525 nm. The fluorescence signal, TTL signals specifying the durations of the 465 or 405 nm LED excitations, and TTL signals from behavioral settings were digitized by a data acquisition module (cDAQ-9178, National Instruments, Austin, TX). The group of digitized signals was simultaneously recorded at a sampling frequency of 1000 Hz by a custom-made LabVIEW program (National Instruments). The fluorescent signal was processed offline. First, each 24-ms excitation period with each LED was identified according to the corresponding TTL signal. One averaged signal value was computed for each excitation period, with two frames at both ends excluded. Each value of 465 nm excitation was paired with a temporally adjacent counterpart of 405 nm excitation and a ratio of the former to the latter was calculated, yielding a processed signal in a ratiometric manner at a frame rate of 20 Hz.

### Operant Conditioning

Behavioral training and tests were performed under constant darkness in an aluminum operant chamber (21.6 × 17.6 × 14.0 cm; Med Associates, Fairfax, VT) housed within a sound-attenuating enclosure. The chamber was equipped with two retractable levers (located 2 cm above the floor), one food magazine between the levers on the floor, and a white noise speaker (80 dB) located on the opposite wall. The mice were required to perform a fixed number of actions (lever presses) to attain a reward: one action on a FR1 schedule or five actions on a FR5 schedule. The mice were first trained for 9–13 consecutive sessions (one 60-minute session/day) on the FR1 schedule followed by six sessions on the FR5 schedule. Training on the FR1 schedule was finished when the mice accomplished seven sessions with more than 50 completed trials per session. The entire experimental procedure took 26–32 days, consisting of surgery, recovery, and training. Each trial began with a sound cue that lasted for 5 s, after which the levers were extended. Presses on the lever on the left of the food magazine (reinforced side) were counted, and a reward pellet (20 mg each, Dustless Precision Pellets, Bio-serv, Flemington, NJ) was dispensed to the magazine immediately after the required number of presses was made. The levers were retracted at the same time as the reward delivery. In contrast, presses on the other lever on the right side had no programmed consequence (non-reinforced side). A refractory period of 0.5 s followed each lever press before the lever was re-extended. In the FR5 sessions, the mice occasionally poked into the magazine before making the required number of lever presses; this was called premature magazine entry (PME). In both PMEs and MEs, the timing of entry was defined as the time point when the distance of the animal’s head to the center of the magazine became less than 2.5 cm. A 30-s inter-trial interval followed each food delivery, during which the levers were not presented and the mice consumed the reward. The subsequent trial was automatically initiated after the interval period ended. TTL signals were generated at the timings of the sound cue, lever extension, and lever press and digitized by a data acquisition module (cDAQ-9178, National Instruments, Austin, TX). The TTL signals were simultaneously recorded at a sampling frequency of 1000 Hz by a custom-made LabVIEW program (National Instruments). Both the FR1 and FR5 sessions lasted for 60 min or until the mice received 100 food rewards. Sound cues were not presented for behavior-only recording using another group of mice without GRAB_DA2m_ expression and without optic fiber implantation. The ITI periods were set at 35 or 30 s. To track the moment-to-moment position of the mice, an infrared video camera (ELP 2 Megapixel WEB Camera, OV2710, Ailipu Technology Co., Ltd, Shenzhen, China) was attached to the ceiling of the enclosure. Reflective tapes were attached to the optical fiber protector (1.2 × 1.4 cm) on the head of the mice. The tapes were recorded at a sampling rate of 20 Hz. The mice’s position in each frame was computed offline by a custom-made MATLAB code.

### Histology

The mice were anesthetized with an overdose of ketamine (100 mg/kg) and xylazine (10 mg/kg) and intracardially perfused with 4% paraformaldehyde phosphate-buffer solution and decapitated. For a better identification of the fiber track, the optical fiber was not removed from the brain until more than 12 h after perfusion. Subsequently, the brains were cryoprotected in 20% sucrose overnight, frozen, and cut into 50-μm thick sections on a cryostat (Leica CM3050 S, Leica Biosystems, Wetzlar, Germany). The sections were mounted on silane-coated glass slides (S9226, Matsunami Glass, Osaka, Japan). The GRAB_DA2m_ signals received no further amplification. Fluorescence images were captured by an all-in-one microscope (BZ-X710, Keyence, Osaka, Japan).

### Data Analyses

A processed ratiometric signal trace was first high-pass filtered at about 0.0167 Hz, corresponding to a wavelength of one minute to exclude low-frequency oscillations. We calculated the z-scores of the DA signal using all inter-trial intervals (ITIs) of a session. Specifically, we first calculated the mean of the DA signals during the last half of the ITI period for each trial and then obtained their mean and standard deviation (SD) for z-score calculation. The reason for only using the last half of the ITI period is that the first half of an ITI period may contain a feeding period. The reason for using all ITI periods of a session for z-score calculation is that there was huge trial-by-trial variability in the DA signal among the ITIs within a session. The variation in the SD of the DA signals during the ITIs *within* the first and second half of a session, respectively, was significantly larger than the variation *between* the first and second half of a session (***Extended Data Fig. 3-1H***), indicating that the DA signals during the last half of the ITI periods fluctuated slightly throughout the entire session but heavily fluctuated on an individual trial-by-trial basis. We assume that the large fluctuation among the trials was caused by spontaneous activities of the mice, such as locomotion and grooming, during the ITI period. We confirmed that the DA dip amplitudes calculated with the z-scores were stable during each session (***Extended Data Fig. 4-3***).

To generate the peri-event plots for the sound cue, lever extension, ME, PME, and head entry at the ITI period (***Figures 3E, 4C and 4F; Extended Data Fig. S4-2A***), the signal was aligned to the event onset timing and then binned temporally into blocks of 100 ms. For each trial, the binned signal was subtracted by the average signal value of the baseline period (–1 to 0 s relative to the sound cue, 0–0.2 s following the ME/PME/entry at ITI).

The response amplitude to an event (***Figures 3F****‒****3H and 4D****‒****4G; Extended Data Figs. S4-1I, S4-2A****‒****S4-2C, S4-3A and S4-3B***) was defined as the mean value of the binned signal during the target period (0–1 s following the sound cue/lever extension, 0.5 to 1.5 s following the ME /PME/entry at ITI) subtracted by that during the baseline period (–1 to 0 s relative to the sound cue, 0–0.2 s following the ME/PME/entry at ITI).

To calculate pre-dip and dip-trough DA levels in ***Figure 4J and 4K***, the mean values of the binned signal at 0–0.2 s and 0.5–1.5 s following PMEs, respectively, were subtracted by that at the ITI baseline period prior to each trial onset (–15 to 0 s relative to the sound cue). Each pre-dip or dip-trough level corresponds to the difference in the Z-score between those during the target window (0–0.2 s or 0.5–1.5 s, respectively) and the corresponding ITI period prior to each trial. The difference between each pair of pre-dip and dip-trough levels therefore corresponds to the DA response size upon PMEs (DA dip amplitude).

### Explained Variance

To compare the contributions of the PME presence, session number, and press number to the inter-press-intervals in each FR5 session, we computed the proportions of the variance explained for each factor. For example, the explained variance of the PME presence was calculated as follows:

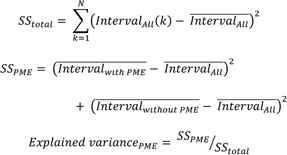

where SS_total_ is the sum of the squared differences between each inter-press-interval and the mean of all inter-press intervals; *Intervals_All_*(*k*)indicates the duration of the k-th inter-press interval of all N intervals in a given session; and 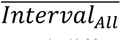 is the average of all interval durations. SS_PME_ specifies the sum of the squared differences between the mean of each condition 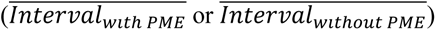 and the mean of all inter-press-intervals.

### PME Percentage, Overall PME Percentage, and PME Bias

To quantify and compare the number of mice’s PMEs across the FR5 sessions, we first computed a value called PME percentage assigned for each inter-press interval group in a session. For example, the PME percentage for the Interval_1–2_, pPME(Interval_1–2_), was calculated as:

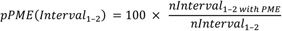

where nInterval_1–2_ refers to the number of inter-press intervals grouped into Interval_1–2_; nInterval_1–2 with PME_ indicates the number of these intervals with a PME. The other PME percentage values, pPME(Interval_2–3_), pPME(Interval_3–4_) and pPME(Interval_4–5_), were calculated likewise.

We also computed the average of these PME percentage values called overall PME percentage assigned for each session. It was calculated as:

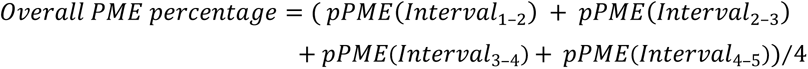

To examine the effects of increasing expectations nearer task completion on the mice making PMEs, we then computed a measure called PME bias for each session on the FR5 schedule. It was calculated as:

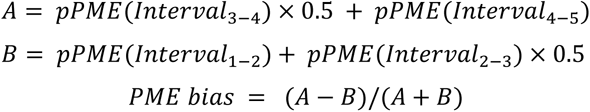

We assigned a weight of 0.5 to data for inter-press intervals located at the middle of the task trial (Interval_2–3_ and Interval_3–4_) by taking their temporal order into account (e.g. in group A above, having a PME during Interval_4–5_ makes the PME timing distribution more biased toward the task completion than having one during Interval_3–4_). The PME bias is given on a scale from –1 to 1. It is close to 1 (–1) if greater expectations make the mice perform more (fewer) PMEs when they are nearer to task completion (i.e., the number of PMEs during Interval_3–4_ or Interval_4–5_ is larger [smaller] than that during Interval_1–2_ or Interval_2–3_). The PME bias is close to 0 if the expectations for task completion have no effect on the number of PMEs (i.e., similar numbers of PMEs are observed during any inter-press intervals).

### Simulation of Behavior and Dopamine Dip

At step s, the temporal difference error was computed, and the values were updated as follows:

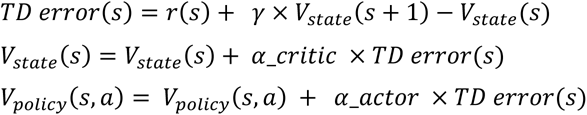

where γ corresponds to the discount factor, and α_critic and α_actor are the learning rates with the state and action, respectively. Suitable parameter values for each mouse’ behavior result were determined via a grid search (α_critic = [0.0001, 0.001, 0.01, 0.1], α_actor = [0.0001, 0.001, 0.01, 0.1], γ = [0.6, 0.7, 0.8, 0.9, 1]). The initial state value and action values (press or PME) were chosen from [0, 1, 2, 3, 4, 5, 6, 7, 8, 9], respectively.

After each lever press, the agents selected their next action—either a press or a PME— according to the mice’s actual selections. We computed a simulated action selection probability for each decision point, taking the probability of making a PME_3‒4_ as an example, as follows:

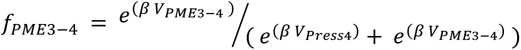

where *β* corresponds to the inverse temperature (chosen from [0.25, 0.5, 1, 2, 4]), and *V _PME3‒4_* and *V _Press4_* correspond to the action values for a PME and a press in inter-press interval group 3‒4 in a given trial, respectively.

In terms of state transition, we introduced sigma values (chosen from [0, 0.25, 0.5, 1, 2]) for forward and backward transitions. When a lever press was selected, the agents jumps 1, 2, 3, 4, or 5 grids forward according to the probability distribution based on the Gaussian kernel (sigma value = original sigma value × number of completed presses in a given trial). The agents were unable to move further beyond the 11^th^ grid—located at the rightmost of our grid world. Any attempt to press at the rightmost grid took the agents to the same 11^th^ grid after the action. Following a PME at the upper row in the grid world, the mice returned (downward shift) to the lower row corresponding to the lever zone. The mice occasionally jumped backward to one of the lever zone grids located at the lower left of the agents’ current position according to the distribution of the gaussian kernel (sigma value = original sigma value × number of completed presses in a given trial). The agents were unable to move further beyond the start grid located at the leftmost of our grid world. Any such attempt took the agents to the starting zone. We started the fitting process from the initial trial of the first FR5 session in each mouse. After the first session, the fitted parameter values were taken over to the following sessions, and we finished the fitting process at the last trial of the selected session.

The log likelihood (LLH) of a model was computed as follows.

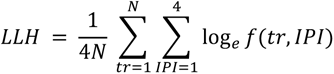

Where *N* corresponds to the number of completed trials in the selected FR5 session of each mouse; *tr* stands for the trial number; and *IPI* stands for the inter-press interval group number. *f(tr, IPI)* represents the probability of the action a mouse selects either a press or PME for an inter-press interval group, *IPI*, in trial *tr*. We defined the best-fitted model as the model with the greatest LLH for each mouse. The non-reinforced lever presses were not taken into account, as the percentage of these was at a negligible level (***Fig. 1F***).

**Figure 1.**
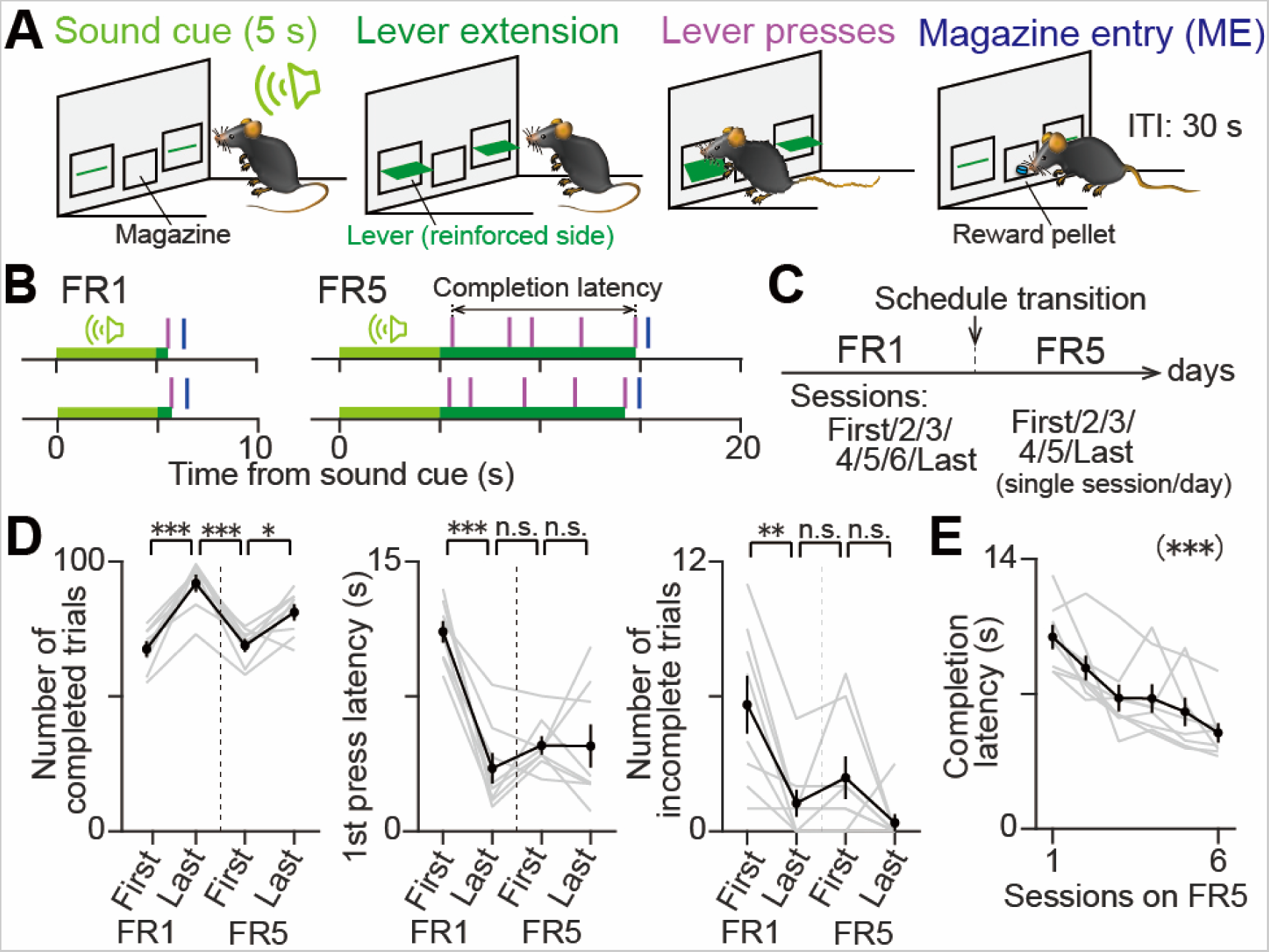
Behavior adjustment across FR5 sessions manifested in completion latency. (A) Procedure of a fixed ratio lever press task. Only the left lever was reinforced; pressing the left lever for a predetermined number of times triggered the delivery of a food reward. If 60 seconds had elapsed from the lever extension onset without the pressing of the reinforced lever enough times, the levers were retracted, and the ITI started without the delivery of a reward. (B) Representative behavior patterns of individual trials on the FR1 (left column) or FR5 (right column) schedule that required pressing the left lever once or five times for a reward, respectively. In each panel, the time axis was aligned with the sound cue onset. Light green bar, sound cue period (0–5 s). Dark green bar, lever extension period (starting 5 s after sound cue onset until the pressing of the last lever required for reward delivery). Vertical magenta tick, lever press. Vertical blue tick, magazine entry (ME). (C) Mice were first trained on the FR1 schedule. Seven consecutive FR1 sessions, each including more than 50 completed trials, were designated as FR1-First, FR1-2, FR1-3, etc. After the FR1 sessions, six FR5 sessions followed. Vertical dashed line, schedule transition from FR1 to FR5. (D) Session-by-session comparisons: number of completed trials (left), first lever press latency (center), and number of uncompleted trials per session (right). It is evident that the association between the stimulus (the lever press) and outcome (the reward) becomes stronger during the FR1 sessions (n = 4 sessions with 8 mice each, Tukey-Kramer post-hoc analysis, FR1-First versus FR1-Last, p = 4.2 × 10^−6^ [number of completed trials], p = 2.0 × 10^−6^ [first press latency], p = 6.2 × 10^−3^ [number of incomplete trials]). While the number of completed trials transiently decreased with the schedule transition from FR1 to FR5 (vertical dotted lines in Fig. 1D), the first press latency and the number of incomplete trials failed to detect behavior adjustment during the FR5 sessions (FR1-Last versus FR5-First, p = 9.6 × 10^−6^ [number of completed trials], p = 0.62 [first press latency], p = 0.77 [number of incomplete trials]; FR5-First versus FR5-Last, p = 1.1 × 10^−2^ [number of completed trials], p > 0.99 [first press latency], p = 0.34 [number of incomplete trials]). * p < 0.05. ** p < 10^−2^. *** p < 10^−5^. (E) Completion latency, the time from the first to last lever press (Fig. 1B, right), detected behavior adjustment during the FR5 sessions (n = 6 sessions with 8 mice each, one-way repeated measures ANOVA, *F*_(5,35)_ = 13.18, p = 3.0 × 10^−7^). Gray line, individual mouse. Black line, mean ± SEM. *** p < 10^−6^.

To check whether the fitting procedure gives reasonable parameter values, we performed the parameter recovery attempt and computed cross-parameter correlations. First, we obtained synthetic behavior data with the initial parameter values pre-defined (α_critic = [0.001, 0.01, 0.1], α_actor = [0.001, 0.01, 0.1], *β* = [0.25, 1, 4], sigma for forward or backward transition = [0, 0.25, 2], initial state value and action values (press or PME) = [0, 4, 8]). 100 parameter sets were randomly selected and used for data synthesis. Next, we fitted the model in the same way as the mice’s behavior data described above using the LLH. Spearman rank correlation coefficients were calculated between the simulated and recovered values for each parameter to examine whether the parameter could be technically recovered within the RL model. Cross parameter correlations between two different parameters were computed using recovered values. If the p-value for a given correlation turned out to be at a non-significant level (corrected p = 0.00078), the corresponding correlation coefficient was regarded as zero.

### Statistical Analyses

We analyzed all data using custom codes written in MATLAB and showed the data as the mean ± standard error of the mean (SEM) unless otherwise described. Sample sizes, which were not predetermined but are similar to our previous reports (Natsubori et al., 2017; Yoshida et al., 2019), are reported in the Results section and figure legends. To determine the statistical significance, we performed one-sample *t*-tests for a single data set (***Extended Data Figs. 5-1E***), one-sample t-tests with Bonferroni corrections for individual data in two or more sample data sets (***Extended Data Fig. 3B***), paired t-tests for two-sample data sets (***Extended Data Figs. 2-2C, 2-2D, 2-2G, 4-1B, and 4-3A***), paired t-tests with Bonferroni corrections for multiple two-sample data sets (***Extended Data Figs. 2-2B, 4-1K, 4-3B, and 4-3C***), and Tukey-Kramer post-hoc analyses for three or more sample data sets (***Figs. 1D, 2E, 3F, 3H, 4D, 4E, Extended Data Figs. 1B, 3H, and 4-1L***). We also employed Kolmogorov-Smirnov tests (***Fig. 2D, Extended Data Figs. 3-1D, 3-1E, 3-1F, and 3-1G***) and Wilcoxon signed rank-sum tests (***Fig. 5F, Extended Data Figs. 4-1E, and 4-1G***) to compare one or two data distributions. We evaluated the statistical significance of variance among four or more sample data sets using one-way repeated-measures ANOVA (***Figs. 1E, 2H, 2I, 3G, 4G, 4J, 4K, 5I, Extended Data Figs. 1-1A, 1-1C, 2-1B, 2-1C, 2-1D, 2-2A, 2-2E, 4-1I, 4-1J, 4-1K, 4-3D, 4-3D, 4-4E, 4-4F, 4-4G, and 5F),*** one-way factorial ANOVA (***Figs. 4F and Extended Data Fig. 4-2A***), and two-way factorial ANOVA (***Figs. 2G, 5H, and Extended Data Fig. 2-1G***). We computed Pearson correlation coefficients to measure linear correlations between the two parameters (***Figs. 4L, 5J, Extended Data Figs. 2-2F, 4-4A, and 4-4C***). All our conclusions in this study are based on the statistical analyses on group data, not by those on individual mouse data. We present results of statistical analyses on single subjects in such as ***Extended Data Figs. 4-2A*** to demonstrate that the effect size is sufficiently large within each subject. The null hypothesis was rejected when p < 0.05.

**Figure 2.**
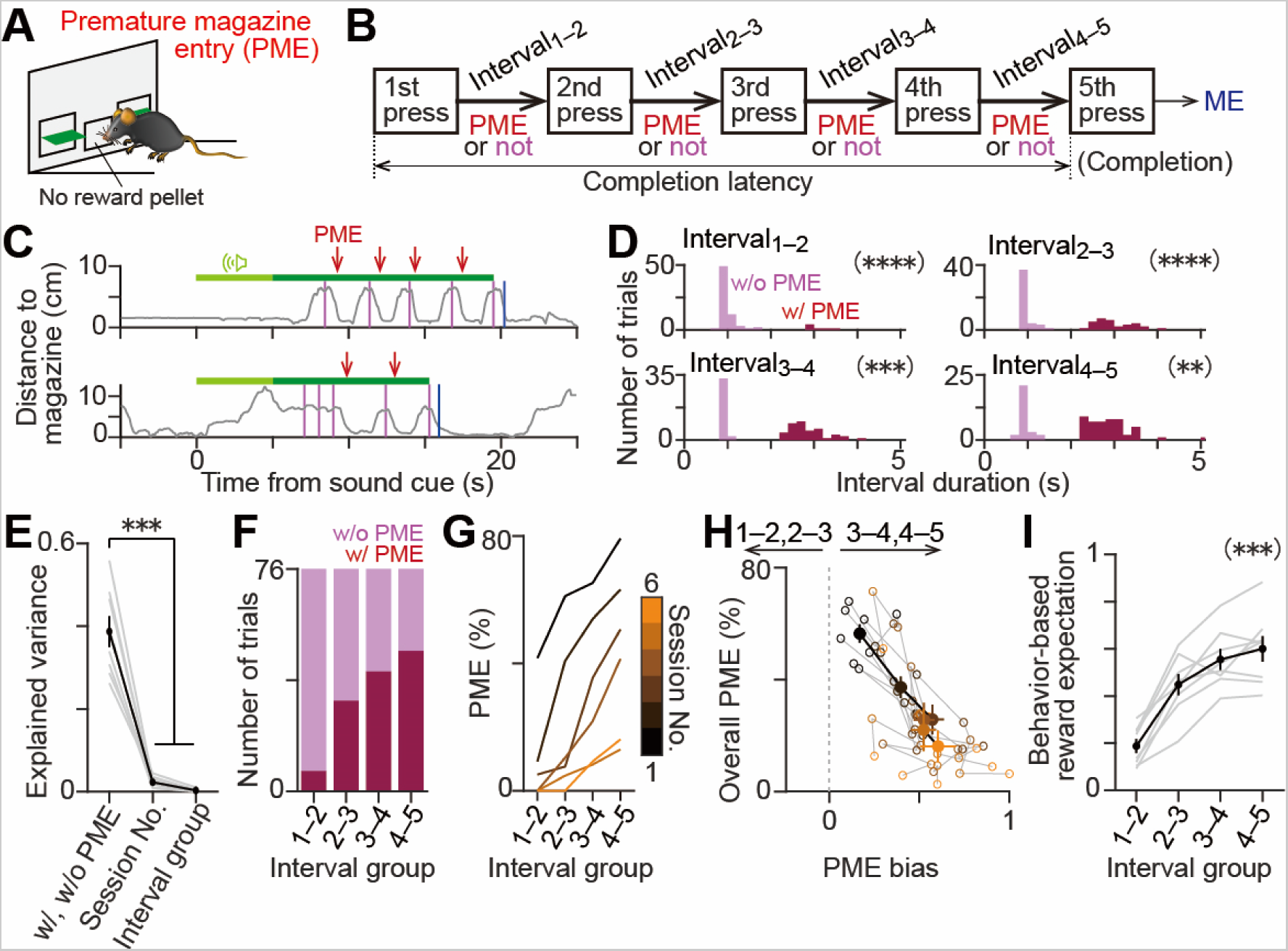
Rising expectations nearer task completion can be quantified by the percentage of a premature magazine entry. (A) The mice occasionally checked the food magazine without a reward before completing the FR5 task, which we called a premature magazine entry (PME). (B) Scheme of the FR5 lever press task. An inter-press interval, a period between two consecutive lever presses, was distinguished by the order of a lever press, e.g., Interval_1–2_ is the period between the first and second lever presses. (C) Representative behavior patterns on the FR5 schedule with PMEs (red arrows) from mouse 9. Each PME was detected based on the distance from the magazine to the mouse’s head. Light green bar, sound cue period. Dark green bar, lever extension period. Vertical magenta tick, lever press. Vertical blue tick, ME. (D) Representative histograms of inter-press interval durations, which were grouped into Interval_1–2_, Interval_2–3_, Interval_3–4_, or Interval_4–5_, with (dark red) or without (pale red) PME in a session of mouse 9. The histogram of inter-press interval durations with a PME was significantly longer than that without a PME (Interval_1–2_, n = 30 and 42 events with and without PME, respectively, Kolmogorov-Smirnov test, p = 6.1 × 10^−16^; Interval_2–3_, n = 44 and 28 events, p = 2.3 × 10^−16^; Interval_3–4_, n = 47 and 25 events, p = 1.2 × 10^−15^; Interval_4–5_, n = 57 and 15 events, p = 1.1 × 10^−11^). ** p < 10^−10^. *** p < 10^−14^. **** p < 10^−15^. (E) Explained variance analysis revealed that the variability of the inter-press interval durations was explained more by the presence or absence of a PME than by the session number (First/2/3/4/5/Last sessions) or inter-press interval group number (Interval_1–2/2–3/3–4/4–5_) (n = 3 factors with 8 mice each, Tukey-Kramer post-hoc analysis, PME versus session number, p = 4.3 × 10^−8^; PME versus inter-press interval group number, p = 2.3 × 10^−8^). Gray line, individual mouse. Black line, mean ± SEM. *** p < 10^−7^. (F) Representative stacked bar chart of trials with (dark red) or without (pale red) a PME during the FR5-2 session of mouse 9. The mouse exhibited more PMEs as the FR5 task completion neared (e.g., the number of trials with a PME during Interval_4–5_ was larger than that during Interval_1–2_). (G) Representative session-by-session plot of the PME percentage versus the inter-press interval groups during the FR5 schedule of mouse 9. While the mean of the PME percentage decreased as the session number increased (early and late sessions indicated as black and brown lines, respectively), the PME percentage within a session increased as the interval number increased, e.g., the PME percentage during Interval_4–5_ was larger than that during Interval_1–2_ (n = 24 data points, 4 interval groups × 6 sessions, two-way factorial ANOVA, interval number, *F*_(3,23)_ = 18.42, p = 2.7 × 10^−5^, session number, *F*_(5,23)_ = 26.13, p = 6.7 × 10^−7^). (H) Group data plot showing an increase in the PME bias from the first (black) to the last (brown) sessions (n = 6 sessions with 8 mice each, one-way repeated measures ANOVA, *F*_(5,35)_ = 16.63, p = 2.1 × 10^−8^) and a decrease in the overall PME percentage (*F*_(5,35)_ = 17.21, p = 1.4 × 10^−8^) during the FR5 schedule. Open circle, individual session (session numbers are color-coded as in G). Filled circle, mean ± SEM within a session. Gray line, individual mouse. Black line, mean of all mice. (I) Behavior-based reward expectations of mice increased as the FR5 task neared completion (n = 4 interval groups with a single session each from 8 mice, one-way repeated measures ANOVA, *F*_(3,21)_ = 71.25, p = 3.6 × 10^−11^). Gray line, individual session. Black line, mean ± SEM. *** p < 10^−10^.

**Figure 3.**
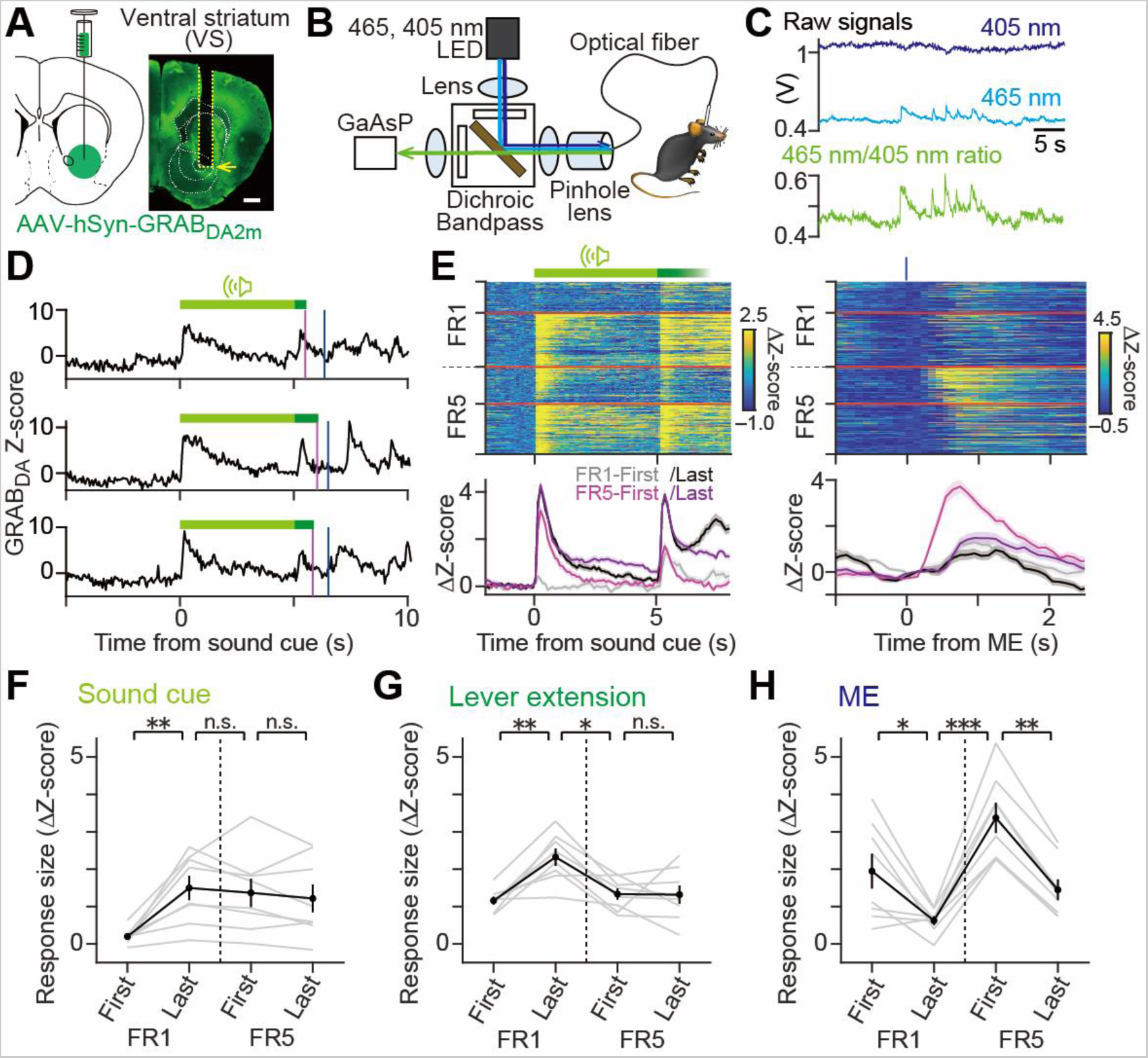
Optical measurement of extracellular DA dynamics in the ventral striatum during a fixed ratio lever press task. (A) Left: Schematic illustration of AAV injection into the VS for the expression of a fluorescent DA sensor, GRAB_DA2m_. Right: Expression pattern of GRAB_DA2m_ (green) in an example mouse and the location of the optic fiber in the VS. Yellow dashed line, optical fiber track. Yellow arrow, fiber tip. Scale bar, 500 μm. (B) Schematic of the fiber photometry system. The light path for fluorescence excitation and emission is through a single multimode fiber connected to the optical fiber implanted in the VS. Excitation light was applied continuously with alternating wavelengths of 405 nm (dark blue arrow) and 465 nm (blue arrow) at 40 Hz. Fluorescence emissions at 525 nm (green arrow) were detected using a GaAsP photomultiplier tube. (C) Example fluorescence intensity fluctuations of GRAB_DA2m_ excited by 465 nm (blue line) or 405 nm (dark blue line) light. Corresponding ratio values (green; 465 nm/405 nm) indicate extracellular DA dynamics. (D) Representative GRAB_DA2m_ signal dynamics during the last session of the FR1 schedule (FR1-Last) of mouse 6. A DA surge was observed after the onset of the sound cue (light green bar) and lever extension (dark green bar). A phasic DA increase also followed an ME (vertical blue line) after a reinforced lever press (vertical magenta line). (E) Top: Representative heatmaps demonstrating GRAB_DA2m_ signal fluctuations during sound cue periods (light green bar in the left panel) followed by lever extension onsets (dark green bar in the left panel) or ME timings (vertical blue line at the top of the right panel) in four selected sessions (from top to bottom, FR1-First, FR1-Last, FR5-First, and FR5-Last sessions). Signals were centered to that from –1 to 0 s at the sound cue onset. The horizontal gray dashed line at the left axis indicates the schedule transition from FR1 to FR5. Bottom: Averaged GRAB_DA2m_ signal traces for respective sessions on the heatmap. (F―H) GRAB_DA2m_ signal dynamics at the lever extension (G) or ME (H) captured the schedule transition from FR1 to FR5 (n = 4 sessions with 8 mice each, Tukey-Kramer post-hoc analysis: FR1-Last versus FR5-First, p = 2.6 × 10^−3^ [lever extension], p = 2.3 × 10^−7^ [ME]), while that at the sound cue onset (F) did not (FR1-Last versus FR5-First, p = 0.96 [sound cue]). Gray line, individual mouse. Black line, mean ± SEM. * p < 0.01. ** p < 10^−3^. *** p < 10^−6^.

**Figure 4.**
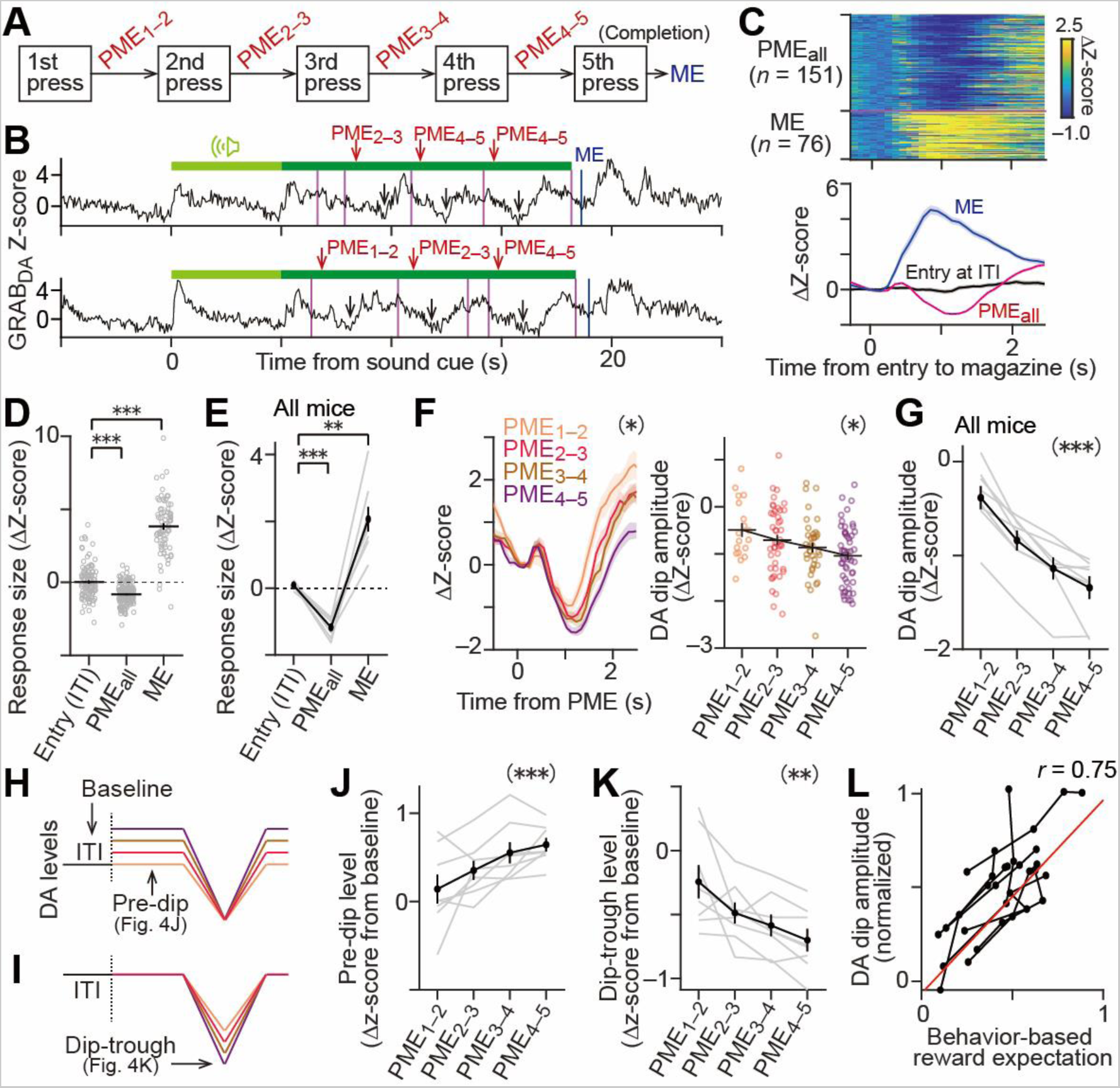
The DA dip amplitude upon PMEs reflects growing reward expectations as completion of FR5 trials approach. (A) Premature magazine entries (PMEs) were grouped by the order of lever presses, e.g., PME_1–2_ is a PME that occurred during the period between the first and second lever presses. ME, magazine entry. (B) Representative GRAB_DA2m_ signal dynamics of mouse 11 during trials with PMEs on the FR5 schedule. A DA dip (black arrow) followed a PME (red arrow). Light and dark green bars, sound cue and lever extension periods. Magenta and blue lines, lever pressing and magazine entry (ME). (C) Top: Representative heatmap of GRAB_DA2m_ signal time courses aligned with the onset of the PME or ME of mouse 11. PME_all_ includes every PME group such as PME_1–2_ and PME_3–4_. Bottom: GRAB_DA2m_ signal traces (mean ± SEM) for PME_all_, ME, and the head entry during ITI on the heatmap with a bin size of 100 ms. A DA dip and surge following the PME (red) and ME (blue), respectively, were evident. GRAB_DA2m_ signal fluctuations during ITI periods were used as the baseline to calculate the z-score. The Δz-score was computed by centralizing the original z-score with that at 0–0.2 s relative to the head entry into the food magazine. (D) Comparison of the DA responses among the ITI, PME, and ME periods of mouse 11 in Fig. 4C. The response size was calculated as the mean Δz-score during 0.5–1.5 s after the head entry in to the magazine (Entry during ITI, n = 109 events, Δz-score = 0.02 ± 0.11; PME_all_, n = 151 events, Δz-score = –0.83 ± 0.06; ME, n = 76 events, Δz-score = 3.82 ± 0.21, Tukey-Kramer post-hoc analysis, entry during ITI versus base (zero), p > 0.99; entry during ITI versus PME_all_, p = 3.8 × 10^−9^; entry during ITI versus ME, p = 4.1 × 10^−9^). Open circle, individual event. Black line, mean ± SEM. *** p < 10^−8^. (E) Group data of DA fluctuations confirmed a significant DA dip and surge after a PME and ME, respectively, when compared to the DA fluctuation upon a magazine entry during an ITI period (n = 8 mice each, Δz-score = 0.08 ± 0.03 [entry during ITI], –1.17 ± 0.1 [PME_all_], 2.07 ± 0.37 [ME], Tukey-Kramer post-hoc analysis, entry during ITI versus base (zero), p = 0.99; entry during ITI versus PME_all_, p = 2.0 × 10^−6^; entry during ITI versus ME, p = 8.0 × 10^−4^). Gray line, individual mouse. Black line, mean ± SEM. ** p < 10^−3^. *** p < 10^−5^. (F) Representative GRAB_DA2m_ traces during the first session of the FR5 schedule (FR5-First) of mouse 11, plotted individually for each PME group. Left: The amplitude of the DA dip following a PME significantly increased when the PME group number increased, e.g., a DA dip following PME_4–5_ was deeper than that following PME_1-2_. Right: Same as the left panel but individual DA dip amplitude values are plotted against PME groups (n = 18–52 events, one-way factorial ANOVA, *F*_(3,150)_ = 3.87, p = 1.1 × 10^−2^). * p < 10^−2^. (G) Group data comparison of DA dip amplitudes among PME groups confirmed deeper DA dips with increasing PME group number (n = 4 PME groups in a single session each from 8 mice, one-way repeated-measures ANOVA, *F*_(3,21)_ = 39.41, p = 8.3 × 10^−9^). Gray line, individual mouse. Black line, mean ± SEM. *** p < 10^−8^. (H, I) Both pre-dip (H) and dip-trough (I) scenarios can realize the larger DA dip size with PME groups. In the pre-dip scenario, modulation of the DA level during the pre-dip period (0–0.2 s from a PME) with a constant DA level during the dip-trough period (0.5–1.5 s after a PME) achieves the larger DA dip size with PME groups. In the dip-trough scenario, the DA level during the pre-dip period is constant but that during the dip-trough period varies to achieve the same outcome for the DA dip size. Line colors correspond to PME group numbers as in Fig. 4F. Note that the z-score was centralized to the baseline during the ITI period prior to each trial. (J) DA levels during the pre-dip period were significantly different among PME group numbers, consistent with the pre-dip scenario (4 PME groups in a session per a mouse from 8 mice each, one-way repeated-measures ANOVA, *F*_(3,21)_ = 8.22, p = 8.3 × 10^−4^). Gray line, individual session. Black line, mean ± SEM. *** p < 10^−3^. Note that z-scores were not centralized to the period at 0–0.2 s relative to the PME, but to the baseline during the ITI period prior to each trial in Fig. 4J, and K. (K) DA levels during a dip-trough period were significantly different among PME group numbers, consistent with the dip-trough scenario (one-way repeated-measures ANOVA, *F*_(3,21)_ =7.54, p = 1.3 × 10^−3^). Gray line, individual session. Black line, mean ± SEM. ** p < 0.01. (L) Variations of the DA dip size upon a PME (a difference in DA levels between pre-dip and dip-trough periods, Fig. 4F) showed a positive correlation with the magnitude of behavior-based reward expectations (Fig. 2I; 4 PME groups in a session per a mouse from 8 mice each; Pearson correlation coefficient, *r* = 0.75, p = 9.2 × 10^−7^). Each dot represents the result of a PME group, such as PME_3–4_, in a session of a mouse. Each line connects results from the same session of a mouse.

**Figure 5.**
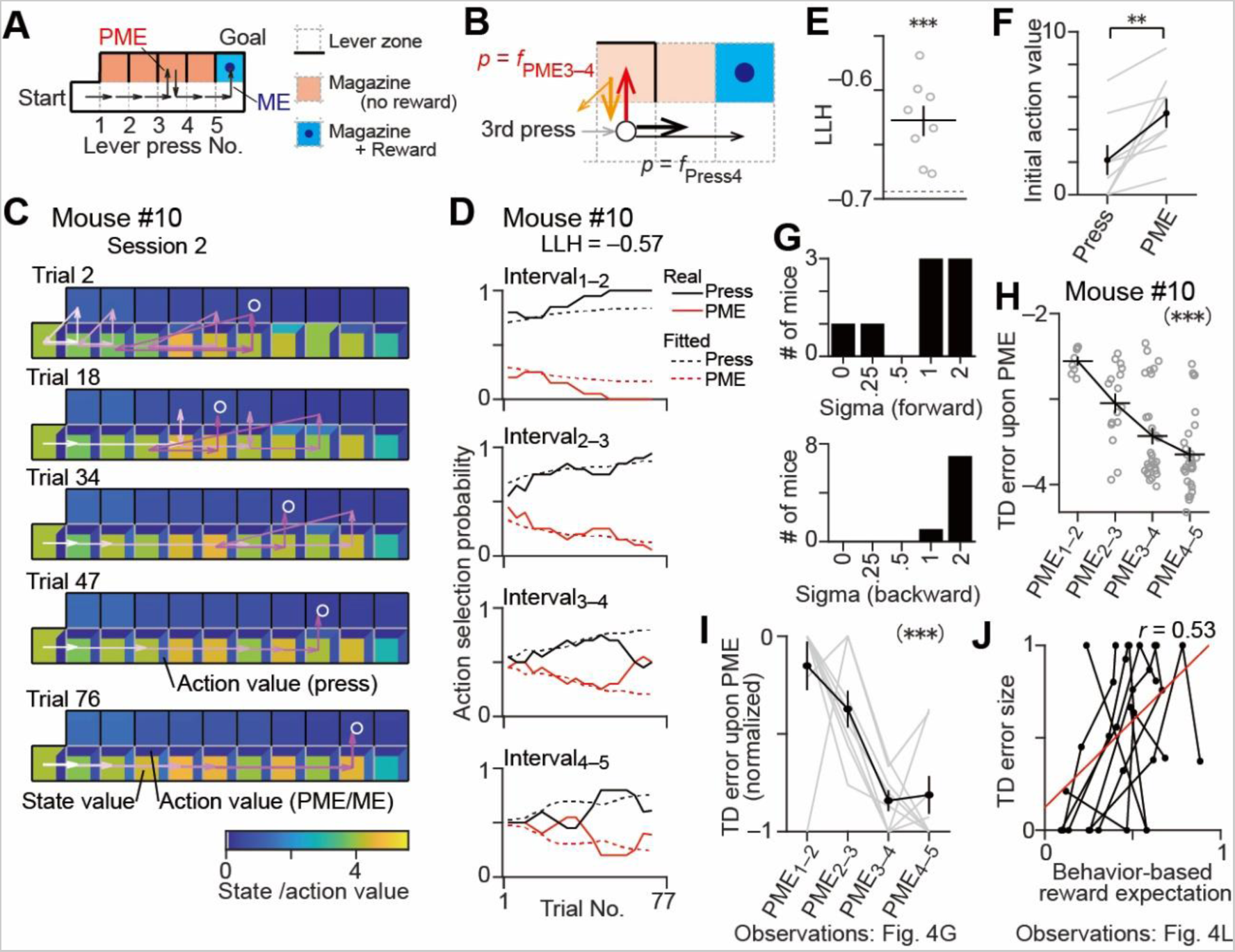
A computational model fitted to the mice’s behavior replicates the increasing trend of DA dips near FR5 task completion. (A) An actor-critic RL model with a TD error in a two-dimensional grid world environment was used to simulate behavioral data of mice in the FR5 schedule. An agent of the model began at the leftmost Start zone, moving step-by-step toward the top right Goal zone for a reward. Horizontal arrow, lever-pressing. Upward arrow, a magazine checking behavior for a PME (red) or ME (blue). (B) A state transition was implemented in the model to simulate occasional misrecognition of the current lever press number by mice. In this example without a state transition, the agent of the model (open circle) chose either a PME_3–4_ (red arrow) with a probability of f_PME3–4_ returning to the same lever zone (thick orange arrow) or the fourth lever press (thick black arrow) with a probability of f_press4_ moving to the next lever zone. With a forward state transition, the agent mistook the fourth lever press with the fifth lever press skipping a lever zone (thin black arrow). With a backward state transition, the agent mistook the third lever press number with the second lever press number (thin orange arrow) after the PME_3–4_. (C) Representative steps (arrows) of the model fitted to the behavioral data of mouse #10. The order of the steps are indicated by graded colors from white (start) to magenta (goal). The state value of the model is indicated by the color of the square in each lever zone. Action values for a PME and a lever press are color-coded at the top and right edges of the lever zone, respectively. Note that the position of the rewarded zone (white open circle) is variable due to the stochastic state transition, demonstrating that the agent does not necessarily follow the lever press number of actual mice behavior during the FR5 schedule. (D) Comparison of time series of action selection probability during completed trials of a representative mouse (solid lines) and the best-fitted model (dashed lines). Action selection probabilities were averaged across 20 trials. (E) Group data of the log likelihood (LLH) per action of the best-fitted model for each mouse. LLHs of the fitted models were significantly larger than that of a random model (dashed line at the bottom) (n = 8 mice, one-sample t-test, LLH = ‒0.63 ± 0.01, *t*_(7)_ = ‒47.47, p = 4.8 × 10^‒10^). *** p < 10^−9^. (F) The initial action value for PMEs was significantly higher than that for the lever presses in the best-fitted models (n = 8 mice, Wilcoxon signed rank test, signed rank = 0, p = 7.8 × 10^‒3^), reflecting the task structure of the preceding FR1 schedule. ** p < 10^−2^. (G) Group data of forward (upper panel) and backward (lower panel) sigma values for state transitions in the best-fitted models. Forward and/or backward sigma values were greater than zero, implying that mice occasionally misrecognized the lever press number. (H) Representative data of the best-fitted model for the behavioral data of Mouse #10, demonstrating that the absolute TD error increased with PME groups (one-way ANOVA, *F*_(3,84)_ = 14.66, p = 1.0 × 10^−7^). Each open circle represents an individual PME event. Black line, mean ± SEM. *** p < 10^−6^. (I) Group data for the normalized TD error amplitude of the best fitted models confirm larger TD errors with PME group numbers (n = 4 PME groups in a single session each from 8 agents, one-way repeated-measures ANOVA, *F*_(3,31)_ = 12.63, p = 2.1 × 10^−5^), reproducing the results of the mice (Fig. 4G). Gray lines, individual data. Black lines, mean ± SEM. *** p < 10^−4^. (J) The best-fitted models showed a positive correlation between the TD error size and the behaviorally estimated reward expectations, reproducing the positive correlation between the DA dip size and the magnitude of reward expectations in experiments (Fig. 4L) (n = 4 PME groups in a single session each from 8 agents; Pearson correlation coefficient, *r* = 0.48, p = 5.3 × 10^−3^). Dots represent each dataset of the best-fitted model. Lines connect data from the same session of the models.

### Data and code availability

The original data (behavior and GRAB signals), source data, and TD model codes are provided on Mendeley Data (http://dx.doi.org/10.17632/4g2yg2cxft.3).

## Results

### Results 1: Completion Latency Decreases with Proficiency in the FR5 Task

We first trained food-restricted mice to press a lever to earn a food pellet. An operant task on a fixed ratio (FR) 1 schedule was used to establish the response (lever press)– outcome (a food pellet delivery) relationship (***Fig. 1A***). In the task, a trial started with a sound cue lasting for 5 s, followed by the presentation of two levers (lever extension). Only the left lever was reinforced; a single press of the reinforced lever resulted in a food reward delivery and the retraction of both levers. Then, the mice poked their head into the food magazine to collect a food pellet (magazine entry, ME). The mice engaged in the series of these events to complete a trial (completed trial). Then, 30 seconds of an inter-trial interval (ITI) was added, followed by a next trial. If the mice failed to press the reinforced lever within 60 s after the lever extension, both levers were retracted and the ITI started without delivering the reward; this was thus considered an incomplete trial. Representative behavior patterns in FR1 trials are shown in ***Fig. 1B***, left side. A single session continued until 100 trials or one hour and was conducted once a day for seven consecutive days (***Fig. 1C***, left side). An FR1-First session was defined as the session in which the mice first accomplished more than 50 completed trials. Establishment of the response–outcome relationship during the seven FR1 sessions was confirmed by an increase in the number of completed trials (***Fig. 1D***, left; FR1-First vs. FR1-Last, Tukey-Kramer, p = 4.2 × 10^−6^). Development of mice’s FR1 task engagement was evident by a decrease in 1) the mice’s reaction time between the lever extension to the first lever press (first press latency; ***Fig. 1D***, center; FR1-First vs. FR1-Last, Tukey-Kramer, p = 2.0 × 10^−6^) and 2) the number of incomplete trials per session (***Fig. 1D***, right; FR1-First vs. FR1-Last, Tukey-Kramer, p = 4.2 × 10^−6^).

After the establishment of the lever press–reward delivery relationship by the seven FR1 sessions, we increased the number of lever presses required for the food reward delivery from one to five. This schedule transition from FR1 to FR5 temporarily decreased the number of completed trials (***Fig. 1D***, left; FR1-Last vs. FR5-First, Tukey-Kramer, p = 9.6 × 10^−6^), but the number was again significantly increased at the end of the FR5 schedule (***Fig. 1D***, left; FR5-First vs. FR5-Last, Tukey-Kramer, p = 1.1 × 10^−2^), implying behavioral adjustment in the mice during the six consecutive FR5 sessions. The behavioral measures for task engagement (first lever press latency and number of incomplete trials) and exploratory behavior (number of non-reinforced lever presses) did not change with the schedule transition (***Fig. 1D***, center and right; Tukey-Kramer, first press latency, p = 0.62; number of incomplete trials, p = 0.77; ***Extended Data Fig. 1-1A***) nor throughout the FR5 sessions (Tukey-Kramer, first press latency, p > 0.99; number of incomplete trials, p = 0.34). These results prompted us to use another behavioral measure to detect the mice’s behavior adjustment during the FR5 schedule.

To quantify the session-by-session adaptation of the mice to the FR5 schedule, we examined the completion latency, defined as the duration from the first lever press to the last (fifth) lever press in the completed trials. A gradual decrease in the completion latency with each FR5 session was observed, indicating that the mice came to perform the lever-press task more smoothly during the FR5 schedule (***Fig. 1E***; one-way repeated measures ANOVA, p = 3.0 × 10^−7^). The percentage of incomplete trials per session was negligible after the mice had established the lever press–reward delivery relationship during the FR1 schedule (1.5 ± 0.8% and 3.5 ± 1.5% of all trials in the FR1-Last and FR5-First sessions, respectively, n = 8 mice). Taken together, these findings showed that 1) once establishing an action–outcome relationship, the mice maintained their degree of engagement even after the ratio requirement was changed from FR1 to FR5 (***Fig. 1D***, center and right), and 2) the mice adjusted themselves to the FR5 sessions to efficiently attain beneficial outcomes by shortening their task completion duration (***Fig. 1E***).

### Results 2: Magazine-Checking Behavior Quantifies Increasing Expectations for Task Completion

We next investigated behavioral factors that accounted for the gradual decrease in the completion latency in FR5 sessions. We noticed that the mice occasionally checked the food magazine without the reward before completing the FR5 task during an inter-press interval between two consecutive lever presses (***Fig. 2A and 2B***). Because 99.0% of the checking behavior consisted of a single head entry to the magazine (4046 out of 4088 total cases from 8 mice), we defined the first head entry during an inter-press interval as a premature magazine entry (PME; red arrows in ***Fig. 2C***). After a PME, mice returned to the lever zone for a next press. We categorized the inter-press intervals into four groups according to the lever press order (e.g., Interval_2–3_ represents the inter-press interval between the second and third lever presses) and examined the effect of the presence of a PME on the length of the inter-press interval, which constitutes the completion latency.

We observed that inter-press intervals with PMEs were significantly longer than those without PMEs in each inter-press interval group (***Fig. 2D***; Kolmogorov-Smirnov test, all p < 10^−10^), implying that a PME delays a task completion. Indeed, variability in the inter-press intervals were explained by the presence or absence of a PME rather than by the session number or inter-press interval group (***Fig. 2E***). Note that the number of head entries during the ITI or reward periods did not change across the FR5 sessions (***Extended Data Fig. 2-1A, B, and C***). Therefore, the decrease in the number of PMEs was critical for the reduction in the completion latency across the FR5 sessions (***Fig. 1E***). We evaluated a behavior adaptation of a mouse to the FR5 schedule by examining PMEs in each inter-press interval group (***Fig. 2F***). The number of trials with a PME increased along with the inter-press interval group, implying that the mice became more expectant of a food reward during later inter-press intervals.

This interpretation assumes that the mice associated multiple lever presses with a reward in the FR5 schedule, instead of just associating lever pressing with a reward. To verify this idea, we compared the number of lever presses the mice made before their first entry to the magazine (either as a first PME or ME) in each trial among the FR5 sessions (***Extended Data Fig. 2-1D***). The average number of lever presses before a first entry to the magazine increased gradually with the FR5 session number, demonstrating that the mice learned to press a lever multiple times for a reward. We also compared the number of lever presses before a first entry to the magazine using the real and shuffled data, in which the interval group number of a PME was shuffled (e.g., a PME of Interval_1–2_ was randomly assigned to a PME of Interval_3–4_). The number of lever presses before a first entry to the magazine in the real data was significantly larger than that of the shuffled (1000 times) data in almost all sessions of all mice (n = 8, ***Extended Data Fig. 2-1E, F***). These results support the idea that mice did not perform a PME randomly but based on their recognition of the lever press number with expectation for a reward.

The percentage of the number of inter-press intervals with PMEs in completed trials (PME percentage) was used as a sensitive measure for a behavior adjustment during the FR5 schedule. The PME percentage decreased as the FR5 schedule proceeded (***Fig. 2G***, early and late sessions indicated by black and brown lines, respectively) in all inter-press interval groups (***Fig. 2G***, x-axis) while maintaining the tendency of a larger PME percentage with later inter-press interval groups such as Interval_3–4_ or Interval_4–5_ (***Fig. 2G***, rising lines; two-way ANOVA, inter-press interval group, p = 2.7 × 10^−5^; session, p = 6.7 × 10^−7^). We quantified whether a PME occurred more frequently in the first half (Interval_1–2_ and Interval_2–3_) or second half (Interval_3–4_ and Interval_4–5_) of the interval groups in each FR5 session by calculating a PME bias; it was closer to –1 or 1 if a PME occurred more frequently in the first or last half of the interval groups, respectively; it was 0 if a PME occurred evenly in the first and second half of the interval groups. Plotting the mean PME percentage across all inter-press interval groups (Overall PME (%)) vs. the PME bias in each FR5 session in each mouse showed that the PME bias was near 0 in the early sessions and gradually shifted toward 1 in the later sessions (from black to brown open circles in ***Fig. 2H***; one-way repeated measures ANOVA, p = 2.1 × 10^−8^), confirming that the overall PME percentage decreased gradually with the FR5 sessions (***Fig. 2H***, y-axis; one-way repeated measures ANOVA, p = 1.4 × 10^−8^).

We selected a single session from each animal that exhibited a major behavior adjustment, which we used for subsequent analysis in this paper (hereafter, such a session will be referred to as the selected FR5 session). The session selection was based on three criteria: 1) the number of PMEs in each inter-press interval group was 7 or more (***Extended Data Fig. 2-2A***), 2) the number of the inter-press interval groups (such as Interval_1–2_ or Interval_2–3_) that accompanied decrease in the number of PMEs between the initial and last 20 trials was the largest among the six FR5 sessions. Typically, the number of the interval groups were three to four in an FR5 session (***Extended Data Fig. 2-2B***), and 3) if there were multiple sessions that satisfied the previous two criteria, the session with the largest reduction rate in the number of PMEs between the initial and last 20 trials was chosen. The selected FR5 sessions reproduced our previous results (***Extended Data Fig. 2-2A vs Fig. 2F***; ***Extended Data Fig. 2-2C vs Fig. 2H***). The selected FR5 session number for each mouse is indicated in ***Extended Data Fig. S4-2***. Mice stayed at the food magazine longer after a ME than a PME, reflecting a consumptive behavior of a mice at the magazine after a ME (***Extended Data Fig. 2-2D***).

Because PMEs constitute a reward-seeking behavior, we estimated a behavior-based reward expectation of a mouse by converting the PME percentage in each inter-press interval group of the selected sessions to a ratio. We compared the relationship between the reward expectation and inter-press interval groups, finding that the behavior based-reward expectations increased with the inter-press interval groups (***Fig. 2I***; one-way repeated measures ANOVA, p = 3.6 × 10^−11^). This result corroborates our interpretation that reward expectation of a mouse increased nearer the task completion.

### Results 3: DA Response upon ME is Modulated by the Schedule Transition from FR1 to FR5

To clarify the DA dynamics during the behavior adjustment, we recorded the extracellular DA fluctuation. We used a viral approach (PHP.eB AAV-hSyn-GRAB-DA2m-W, 1.0 × 10^14^ genome copies/ml) to express a genetically encoded optical dopamine sensor— GRAB_DA2m_—in the ventral striatum (VS) (***Fig. 3A; refer also to Extended Data Fig. 3-1A*** for mediolateral or rostrocaudal variations of the recording sites among all mice used). We targeted the VS because our previous studies demonstrated that the VS played a critical role in lever pressing operant tasks under FR schedules (Natsubori et al., 2017; Tsutsui-Kimura et al., 2017a; Tsutsui-Kimura et al., 2017b; Yoshida et al., 2020). We developed a fiber photometry system to continuously monitor extracellular dopamine level fluctuations *in vivo* through an optical fiber during the operant task (***Fig. 3B***). The GRAB_DA2m_ sensor was alternately excited by 405 nm and 465 nm lasers at 40 Hz, and the emissions were detected by a GaAsP photocathode with a time resolution of 50 ms. The fluorescence intensity ratio (465 nm/405 nm) captured the extracellular DA fluctuations on a timescale of sub-seconds (***Fig. 3C***). The fluorescence intensity was stable for an hour with little photobleaching effects, enabling consistent DA measurements for a task session (***Extended Data Fig. 3-1B, C***). Our fiber photometry system provided a direct measurement of extracellular DA fluctuations at a target site of DA neurons, complementing previous studies that investigated the activity of midbrain DA neurons with electrophysiological, electrochemical, or optical measurements (Amo et al., 2020; Schultz et al., 1997; Starkweather and Uchida, 2021). It has been reported that the expression of GRAB sensors did not change its downstream intracellular signaling (Sun et al., 2018). We confirmed that neither GRAB_DA2m_ expression nor the optic fiber implantation in our study altered mice’s behavior (***Extended Data Fig. 1-1B, C; Extended Data Fig. 2-1B, C***).

We succeeded in observing an extracellular DA level increase at the onset of the sound cue, lever extension, or ME in the FR operant task (***Fig. 3D and Fig. 3E***). These DA dynamics were almost in line with previous studies in three aspects of Pavlovian and operant conditioning (Amo et al., 2020) (Schultz et al., 1997); First, it is reported that there is a large DA response to an unconditioned stimulus (US) and an almost negligible response to a conditioned stimulus (CS) at the beginning of the CS–US association. In our study, there were a clear DA response upon the US (gray trace for DA dynamics upon an ME during FR1-First in the bottom right panel of ***Fig. 3E***) and relatively small DA responses upon the CSs (gray traces for DA fluctuation upon the sound cue and lever extension during FR1-First in the bottom left panel of ***Fig. 3E***) during the FR1-First session. Secondly, it is reported that a DA activity shows clear responses to the CS but no obvious responses to the US once the CS–US association has been established. In our study, there were a significant increase of DA responses upon the CSs (black trace for DA responses upon a sound-cue [green bar] or lever extension [dark green bar] in the bottom left panel in ***Fig. 3E***; group data comparison of the DA response amplitude between FR1-First vs FR1-Last upon a sound cue in ***Fig. 3F*** and a lever extension in ***Fig. 3G***; Tukey-Kramer, p = 3.5 × 10^−4^ [sound cue], p = 4.7 × 10^−4^ [lever extension]), and a significant decrease of a DA response upon the US (a black trace for DA fluctuation upon a ME at bottom right panel in ***Fig. 3E***; group data comparison of the DA response amplitude upon a ME between FR1-First vs FR1-Last in ***Fig. 3H***; Tukey-Kramer, p = 3.1 × 10^−3^) from the FR1-First to FR1-Last sessions. And thirdly, it is reported that DA responses to the US and CS increases and diminishes, respectively, returning to the initial state of the CS– US association upon a violation of the CS–US association. In our study, there were an augmentation of the DA response upon the US (black versus light purple traces upon a ME in the bottom right panel of ***Fig. 3E***; group data comparison of the DA response amplitude upon a ME between FR1-Last vs FR5-First in ***Fig. 3H***, Tukey-Kramer, p = 2.3 × 10^−7^), and a decrease of a DA fluctuation upon the lever extension (black versus light purple traces for DA fluctuation upon a lever extension [dark green bar] at left panels in ***Fig. 3E***; group data comparison of DA response amplitudes upon a lever extension between FR1-Last vs FR5-First in ***Fig. 3G***, Tukey-Kramer, p = 2.6 × 10^−3^) from the FR1-Last to FR5-First sessions.

The above results demonstrate that 1) an association between the food reward and lever pressing was established during the FR1 sessions, and 2) the schedule transition from the FR1-Last to FR5-First sessions violated the CS–US association. A minor difference of DA dynamics between our observation and the previous studies was that the DA response to another CS (sound cue) did not diminish and remained large even after the schedule transition (***Fig. 3F***, FR1-Last versus FR5-First; Tukey-Kramer, p = 0.96). This observation suggests that the schedule transition did not diminish completely the association between the CS–US association, which might explain why the mice maintained their motivation and continued to perform operant responses even after the schedule transition (***Fig. 1D***, 1^st^ press latency and number of incomplete trials between FR1-Last vs FR5-First). Of note, the DA level increase upon lever pressing was not significant throughout our study (magenta lines in ***Fig. 3D***), which was consistent with a previous report that activity of DA axons projecting to the ventral striatum was not as modulated by mice’s motion as that projecting to the dorsal striatum (Howe and Dombeck, 2016).

### Results 4: DA Dip Amplitude Increases with Reward Expectations

We analyzed the DA dynamics during PMEs in detail because the PME ratio reflected the reward expectations of the mice (***Fig. 2I***) and PMEs were the only cue for mice to realize the schedule transition from FR1 to FR5. For later analysis, PMEs were classified into four groups according to the lever press order, e.g., a PME_2–3_ is a PME that occurred between the second and the third lever presses (***Fig. 4A***). We noticed a transient drop in the DA levels (***Fig. 4B and Extended Data Fig. 4-5***, black arrows just above the Z-scored DA fluctuation) following a PME (***Fig. 4B***, red arrows at the top) of a representative mouse. The mean time-course of the DA fluctuation of all trials in the selected session of this mouse demonstrated a transient decrease in the DA level upon a PME (DA dip; ***Fig. 4C*** bottom, red trace; PME_all_ includes all four PME groups), in addition to the DA increase upon an ME (DA surge; ***Fig. 4C*** bottom, blue trace). It is noteworthy that the same head entry behavior during the ITI periods, which was the lowest possible state of reward expectation during the FR5 task, did not evoke such a DA fluctuation (***Fig. 4C***, black trace). We assume that these head entries during the ITI period are different from such goal-directed reward-seeking behavior during the trial period. This is because 1) they have already learned that an operant response(s) is needed for the pellet reward to be delivered, 1) such head entries were made right after the mice have reached a goal and have consumed the corresponding reward, and 3) the number of head entries during ITIs did not change across FR5 sessions (***Extended Data Fig. 2-1B, C***, ITI). Alternatively, such head entries during the ITI period might be the behavior to confirm that there is no fraction of a food pellet accidentally left by the mice. In either case, the degree of reward expectations would be lower than one full food pellet like those during the trial period. Statistical analyses confirmed the above DA dynamics (***Fig. 4D*** for the representative mouse; ***Fig. 4E*** for all mice). These results indicate that the DA fluctuation upon a head entry depends on the magnitude of mice’s reward expectations.

We found that the amplitude of a DA dip within a session increased with the PME groups (***Fig. 4F*** and ***Extended Data Fig. 4-2A*** for DA dip traces of a representative mouse and all the mice, respectively; ***Fig. 4G*** for a group data comparison of the DA dip amplitude), implying that proximity to the completion of the task affected the amplitude of the DA dip. In these analyses, the DA dip amplitude was calculated as a difference in the z-score (Δz-score) from the magazine entry timing, assuming a constant pre-dip DA level (0–0.2 s relative to PMEs) and emphasizing the role of fast DA fluctuations on the timescale of seconds (***Fig. 4I***; a dip-trough scenario). Another possibility for realizing the variations of the DA dip amplitude was a modulation of the tonic DA level preceding a DA dip with a constant DA dip-trough level (0.5–1.5 s relative to PMEs) (***Fig. 4H***; a pre-dip scenario), which emphasized the role of tonic DA fluctuations for achieving a quantitative representation of a negative RPE (Hamid et al., 2016). We assessed the two scenarios by analyzing DA level variations at pre-dip and dip-trough periods (***Fig. 4H, I***); The z-score of the DA signal during the entire session was calculated using the ITI period as the baseline because the DA level during the ITI period was stable among the PME groups (***Extended Data Fig. 4-4G***). Each pre-dip or dip-trough level corresponds to the difference in the Z-score between those during such a target window and the corresponding ITI period prior to each trial. We revealed that both scenarios were valid in our study. DA levels both at pre-dip and dip-trough periods showed increasing and decreasing trends, respectively, with PME groups (***Fig. 4J, K***). Thus, increase of pre-dip level and decrease of dip-trough level of DA fluctuations sculpted the larger DA dip amplitudes with the mice’s increasing proximity to task completion.

We discovered a significant positive correlation between the amplitude of the DA dip upon a PME and the magnitude of behavior-based reward expectations (***Fig. 4L***). We hypothesized that the DA dip size upon a PME quantified the magnitude of mice’s reward expectations because 1) the RL theory formulates the RPE as the difference in value between the achieved reward and expected reward, 2) DA dynamics reflect the RPE, and 3) the achieved reward after the PME was zero (no reward) in the current study. It is noteworthy that the DA surge upon a FR5 task completion was unlikely to be involved in behavioral refinement by reducing the number of PMEs because the amplitude of the DA surge upon an ME did not distinguish the number of PMEs the mice made in the corresponding trial (***Extended Data Fig. 4-4F***), indicating that behavior refinement in the FR5 task depends on DA dynamics upon PMEs rather than that upon a task completion (MEs).

The DA dip upon a PME was consistent and reproducible. The z-scored DA signal 0.5‒1.5 s after a PME was below zero in 90.3 ± 2.0% of all PMEs (***Extended Data Fig. 4-1E, F***), and the DA level decreased upon a PME in 89.9 ± 1.4% of all PMEs (***Extended Data Fig. 4-1G, H***). The time interval of 0.5‒1.5 s after a PME was selected because the negative peak of the DA dip occurred 1.09 ± 0.03 s after a PME on average (n = 8 mice, ***Extended Data Fig. 4-1C, D***). This delay includes approaching behavior to the magazine because a PME was detected when mice came within 2.5 cm of the magazine. Therefore, we analyzed the locomotive speed of mice upon a PME and found a stereotypical behavior of mice that they arrived at and started to explore the magazine 0.37 ± 0.02 s after the PME detection (***Extended Data Fig. 4-1A, B***). The frequency of PMEs per session decreased with FR5 sessions (***Extended Data Fig. 2-1B, C***, IPI; ***Extended Data Fig. 2-1G***), just as that in the mean amplitude of the DA surge upon an ME (***Extended Data Fig. 4-1I***, right), corresponding to the establishment of the CS– US association (***Fig. 1E***). Although the average amplitude of all DA dips did not change with the sessions (***Extended Data Fig. 4-1I***, left), classifying the DA dips according to the PME group number revealed that 1) the DA dip amplitude upon the PME_1-2_, PME_2-3_, or PME_3-4_ group is greatest in the first FR5 session, and that 2) there is a trend of the progressive DA amplitude reduction with training in the earlier PME group numbers (DA dips upon PME_1-2_, PME_2-3_) in accordance with the behavior results (***Extended Data Fig. 4-2B and 4-2C***). These results imply that the relationship between the DA dip amplitude and the behavior-based reward expectations we observed within a session may hold across the task sessions. The frequencies per minute of PMEs (paired with a DA dip) and MEs (paired with a DA surge) were 1.9 ± 0.1 and 1.2 ± 0.03, respectively, in the selected FR5 sessions (***Extended Data Fig. 4-1J, K, L***). The frequencies per trial of PMEs were 1.8 ± 0.2 in the selected FR5 sessions.

We considered two possibilities against our interpretation that the amplitude of a DA dip increased with PME groups. The first one assumes the following two cases: 1) the amplitude of a DA dip increased (or decreased) with trials within a session regardless of the corresponding PME group, and 2) only PMEs in larger PME groups appeared more in later (or earlier) trials. Combination of these two situations would result in an apparent increase in the DA dip amplitude with PME groups even if there is no relation between the DA-dip depth and PME group. To test this possibility, we compared the DA dip amplitude in the first and the second half of a session and revealed no difference in DA dip amplitude between the two halves (***Extended Data Fig. 4-3A, B***). In addition, the number of PMEs showed a decreasing trend within a session regardless of the PME groups (***Extended Data Fig. 4-3C, D***). These results did not support the first possibility. The second possibility is related to a waiting time from the mice’s task engagement (the first lever press in a trial) to a PME. If there is a positive correlation between the waiting time and the amplitude of a DA dip, DA dip amplitudes would increase with PME groups even if the DA dip amplitude and PME group were unrelated. To test this possibility, we calculated a correlation coefficient between the waiting time and DA dip amplitude upon a PME (***Extended Data Fig. 4-4A, B***). Note that the correlation coefficient was calculated for each PME group and that PME groups were further divided based on the number of preceding PMEs in the same trial, e.g., PME_3‒4 (0)_ and PME_3‒4 (1)_ were PME_3‒4_ without and with a preceding PME, respectively. These subdivisions were necessary to subtract the effects of the number of presses and PMEs on the waiting time because the waiting time naturally increases with the press number (***Extended Data Fig. 2-2F***) or the presence of a PME (***Extended Data Fig. 2-2G; Fig. 2D***). After these subdivisions, we found no correlation between the waiting time and DA dip amplitude in more than 95% of the cases (***Extended Data Fig. 4-4A and S4-4B***). These results did not support the second possibility. Therefore, we concluded that the DA dip amplitude increased with PME groups.

### Results 5: RL Model Reproduced Behavioral Adjustment and DA Dip Amplitude Variations

We employed computational modeling of behavioral data to address whether the DA dip size variations had any functional role in behavior adjustment during the FR5 schedule. We used an actor-critic RL model with temporal difference (TD) errors because DA fluctuations have been proposed to represent TD errors, which serve as teaching signals to guide experience-based behavior adjustment (Amo et al., 2020; Moutoussis et al., 2009; O’Doherty et al., 2003; Schultz et al., 1997; Starkweather and Uchida, 2021; Suri and Schultz, 2001). The agent in the model has a state value and action values for a forward step (lever-pressing) and an upward step (checking a magazine: ME or PME) at each lever zone in a two-dimensional grid world environment (***Fig. 5A***). The agent begins at the leftmost start zone in the environment and chooses an action for a forward or upward step according to the softmax action selection rule based on the action values (Materials and Methods). The agent updates the state and action values step-by-step using a state value-derived TD error. We assumed that the mice might occasionally misrecognize the current lever press number, e.g., mice might mistake the second lever press for the fourth one, because there was no external cue about the number. To introduce this stochastic error made by the mice, we implemented a non-deterministic “state transition” (Starkweather et al., 2018) in our model; Even if the agent chooses an action for a single forward step, the agent moves 1 to a maximum of 5 forward step(s) in the grid world (***Fig. 5B***, thick and thin black arrows, respectively). Similarly, the agent’s choice of an action for a PME is followed by zero to a maximum of 5 backward step(s) (***Fig. 5B***, thick and thin orange arrows, respectively). We performed a grid search to fit eight free parameters of the model for the behavioral data of the mice during the FR5 schedule (***Fig. 5C***). The free parameters were the initial state value for a lever zone, the initial action values for a lever press and a PME, learning rates for state and action values, an inverse temperature for the softmax function, and forward and backward sigma values that specified the probabilistic state transition. Goodness of a fit was assessed by calculating a log likelihood (LLH) of action selection probabilities of a model for a lever press and a PME (***Fig. 5D***, comparison of observed and fitted action selection probabilities). To check whether the fitting procedure using our RL model gives reasonable parameter values, we checked parameter recoveries and cross-parameter correlations. We first obtained synthetic behavior data with the pre-defined initial parameter values and then fitted the model to such synthesized data in the same way we performed for the mice’s actual behavior data. All parameters were recovered with significant correlation coefficients (Extended Data Fig. 5A, p ranging from 2.5 × 10^‒20^ to 5.8 × 10^‒4^, Spearman rank correlation coefficients from 0.34 to 0.76). Furthermore, the correlation between each pair of two different parameters was not significant or weaker (Extended Data Fig. 5B). These results show that our parameter optimization procedure for the RL model is capable of recovering the original parameters.

We found that the best fitted models had significantly higher LLH values than a random model that chose a lever press or a PME with an equal probability (***Fig. 5E***; one-sample t-test, LLH = ‒0.63 ± 0.01, p = 4.8 × 10^‒10^). The best fitted models also had significantly higher initial action values for a PME than that for a lever press (***Fig. 5F***; Wilcoxon signed rank test, p = 7.8 × 10^‒3^), suggesting that the CS–US association during the FR1 schedule biased action values for a PME over a lever press during initial sessions of the FR5 schedule. The best-fitted models had sigma values larger than zero for the state transition, implying that mice actually misrecognized the current lever press number. We discovered that absolute TD errors of the best-fitted models increased with PME group numbers (***Fig. 5H***, representative data; ***Fig. 5I***, group data), reproducing the larger DA dip amplitudes of the mice with PME group numbers (***Fig. 4F, G***). Moreover, the best-fitted models showed a positive correlation between the TD error size and the magnitude of reward expectations estimated based on the PME percentage (***Fig. 5J***), corresponding to the experimental data (***Fig. 4L***). These results with computational modeling supported the functional relevance of the DA dip amplitude variations for the behavior adjustment during the FR5 schedule.

## Discussion

We addressed the neuronal representation of the magnitude of reward expectations. Our data from behavioral experiments, DA fluorescence observations, and computational modeling of behavioral data consistently supported the interpretation that the magnitude of the DA dip is associated with the magnitude of reward expectations. This interpretation is in line with previous reports that dopaminergic neuronal firing is reduced upon the omission of an expected reward (Schultz et al., 1997) and the DA level decreased transiently upon head entries to the reward location without a reward during an operant task in a progressive ratio (PR) schedule (Ko and Wanat, 2016).

The simple operant task in this study had three advantages for our finding that the DA dip amplitude reflected the magnitude of reward expectations: 1) the fixed reward size, 2) the stereotypical behavior for a reward expectation, and 3) the modulation of reward expectations in single trials. The reward size was always stable: a single pellet after each FR5 task completion. In other behavioral tasks, the reward size is commonly varied to modulate the magnitude of expectations, e.g., the amount of juice, which might introduce extra DA fluctuations unrelated to reward expectations. PMEs in the FR5 task accompanied an almost identical behavior pattern for reward expectations of different magnitude. This situation contrasts with other tasks in which mice must choose distinct behaviors such as pressing left or right levers for a reward with different expectancy, which might cause extra DA fluctuations reflecting the behavior differences. The magnitude of reward expectations in our FR5 task increased monotonically between the PME groups in a single trial, which was a unique task design when compared to other common tasks such as a lottery economic task, a random-ratio (RR) task, and a progressive ratio (PR) task. Although the latter tasks enable a flexible manipulation of reward expectations by an experimenter, multiple trials with or without rewards are necessary, which might evoke a long timescale DA level fluctuation beyond a single trial, complicating the interpretation of DA fluctuations.

There are two hypotheses for realizing the variations in the DA dip amplitude: tonic and phasic. The first hypothesis assumes that tonic DA modulation realizes variations in the DA dip amplitude (Hamid et al., 2016); A larger DA dip is apparently realized by a higher DA level prior to the dip while the DA concentration during the dip-trough is constant. Another hypothesis proposes that the phasic DA fluctuation modulate the DA dip magnitude (Dabney et al., 2020); A larger DA dip is realized by variable DA levels at the trough while the pre-dip DA level is constant. Our sensitive and stable DA imaging system combined with the simple FR5 operant task was critical to demonstrate that mice’s higher reward expectations upon a PME coincide with a higher pre-dip DA level, and that such a higher pre-dip level is followed by a lower dip-trough DA level, revealing that both tonic and phasic fluctuations of DA levels cooperatively realize the variation in the DA dip amplitude.

The current study on extracellular DA dynamics complements previous works on DA neuron firing for quantitative representation of negative RPE (Hamid et al., 2016; Hart et al., 2014). Early pioneering works focused on firing of DA neurons (Schultz et al., 1997; Tobler et al., 2005), rather than DA itself, to explain variation of the DA dip amplitudes and negative RPEs. Indeed, theoretical works have proved that a pause in the DA neuron firing realized a continuous representation of both positive and negative RPE (Bayer et al., 2007) and a linear modulation in the DA dip amplitudes (Dreyer et al., 2010). Experimental studies confirmed that the larger the reward probability, the larger the number of DA neurons that paused their firing upon omission of the reward (Tian and Uchida, 2015). These results indicate that firing of DA neurons alone can recapture the DA dip amplitude variations. However, unexpected DA release for reward expectations without changes in the firing of DA neurons has been found (Mohebi et al., 2019). Moreover, presynaptic modulation of the striatal DA release by cholinergic (Cachope et al., 2012; Chéramy et al., 1986; Threlfell et al., 2012; Zhou et al., 2001) or glutamatergic (Glowinski et al., 1988) innervation has been reported, implying DA release regulation independent of DA neuron firing. A prior work using FSCV has identified variations in the amplitudes of the surge and dip of the extracellular DA concentration in the rat ventral striatum after a lottery task, demonstrating symmetrical representation of positive and negative RPE, respectively (***Hart et al., 2014***). Ko and Wanat have succeeded in isolating a DA dip in the ventral striatum following a PME in an operant task using a PR schedule with FSCV (***Ko and Wanat, 2016***), but modulation of the DA dip amplitude within a trial had yet to be tested. In the present study, we demonstrated the modulation of the DA dip amplitude by the magnitude of reward expectations. It is also noteworthy that such monotonic variations in such a relative DA response size indeed consist of positive and negative correlations of absolute DA levels at pre-dip and dip-trough timings with the magnitude of mice’s reward expectations before and after unexpected reward omissions, respectively. Our findings thus provide evidence that such apparently different mechanisms (related to arguments regarding tonic or phasic DA fluctuations) can reside together in a single behavioral or dopaminergic recording environment to realize and emphasize the DA dip amplitude variations, helping to reconcile the two different theories. Further investigations are necessary to reveal neurophysiological mechanisms that links DA neuron firing and extracellular tonic/phasic DA dynamics with respect to the DA dip size variations.

One caveat of this study is that, if compared across sessions, the DA dip amplitude does not necessarily correspond to the absolute PME rate in some PME groups. We rather found such a relationship within a task session and in a single trial-based manner, and this is one of the novelties of our study. It remains an open question what mechanisms are behind the long-term dynamics in the DA dip amplitude upon an outcome as a result of spontaneous behavior in an operant task. Additionally, one limitation in our DA measurements is that spatial pattern of the virus mediated GRAB_DA2m_ expression did not necessarily match to that of DA receptors in the striatum. While it has been reported that half of the major DA receptors in the striatum, D_1_ and D_2_ receptors, are distributed in the dendrites and 25% in the spine (***Yung et al., 1995***), subcellular localization of GRAB_DA2m_ is unknown in the current study. Therefore, it is not evident whether neurons expressing dopamine receptors can detect the difference in the extracellular DA levels reported here. Even so, considering the recent report that striatal DA receptor-expressing medium spiny neurons (MSNs) can detect a decrease in DA levels for as short as 0.4 seconds (***Iino et al., 2020***), it is highly likely that MSNs are able to distinguish the small differences in tonic and phasic DA levels. Another caveat in the current study is the stability of the virus expression of GRAB_DA2m_. In this study, DA measurements were completed using the same configuration such as the excitation light intensity and detector gain for each mouse within 3‒4 weeks after the virus injection surgery, indicating stable GRAB_DA2m_ expression at least during our experiments.

We investigated the functional relevance of DA dip amplitude variations in behavior adjustment during the FR5 schedule using a computational modeling of the task using a RL algorithm (Bayer and Glimcher, 2005; Collins and Frank, 2014; Hart et al., 2014; Nakahara et al., 2004; Niv et al., 2007). The amplitude of the negative TD error upon a PME increased with PME groups in our agent of the model, reproducing the increasing DA dip amplitude with PME groups in mice. We implemented a state transition in our agent to introduce possible misunderstanding of lever press numbers by mice. Our best fitted agent of the model for each mouse had non-zero transition values, supporting the idea that the mice actually misunderstood the current lever press number occasionally.

## Conclusion

We propose that the degree of reward expectations upon unexpected reward omissions is represented quantitatively by the magnitude of the DA dip in the ventral striatum and that our RL model suggested that DA dip amplitude variations are critical for the behavioral adjustment during the FR5 schedule. Such DA dip amplitude variations might provide a biological basis for quantitative negative reward prediction errors in the reinforcement learning theory and a neurophysiological clue to elucidate the degree of disappointment.

## Author Contributions

Y.S., N.T., and K.F.T. designed the research. K.F.T. supervised the project. S.Y. created the GRAB_DA2m_ virus. Y.S. and N.T. established the GRAB_DA_ fiber photometry system. Y.S. conducted the fiber photometry experiments, wrote the main manuscript text, and prepared the figures. N.T. edited the main manuscript text.

## Acknowledgements

This work was funded by Grant-in-Aid for Brain Mapping by Integrated Neurotechnologies for Disease Studies (Brain/MINDS) JP21dm0207069 from the Agency for Medical Research and Development (AMED), Japan (K.F.T. and S.Y.); JSPS KAKENHI Grant Numbers 19K06944, 21H00212 (N.T.) and 19J01068 (Y.S.).

---

**Extended Data Figure 1-1.**
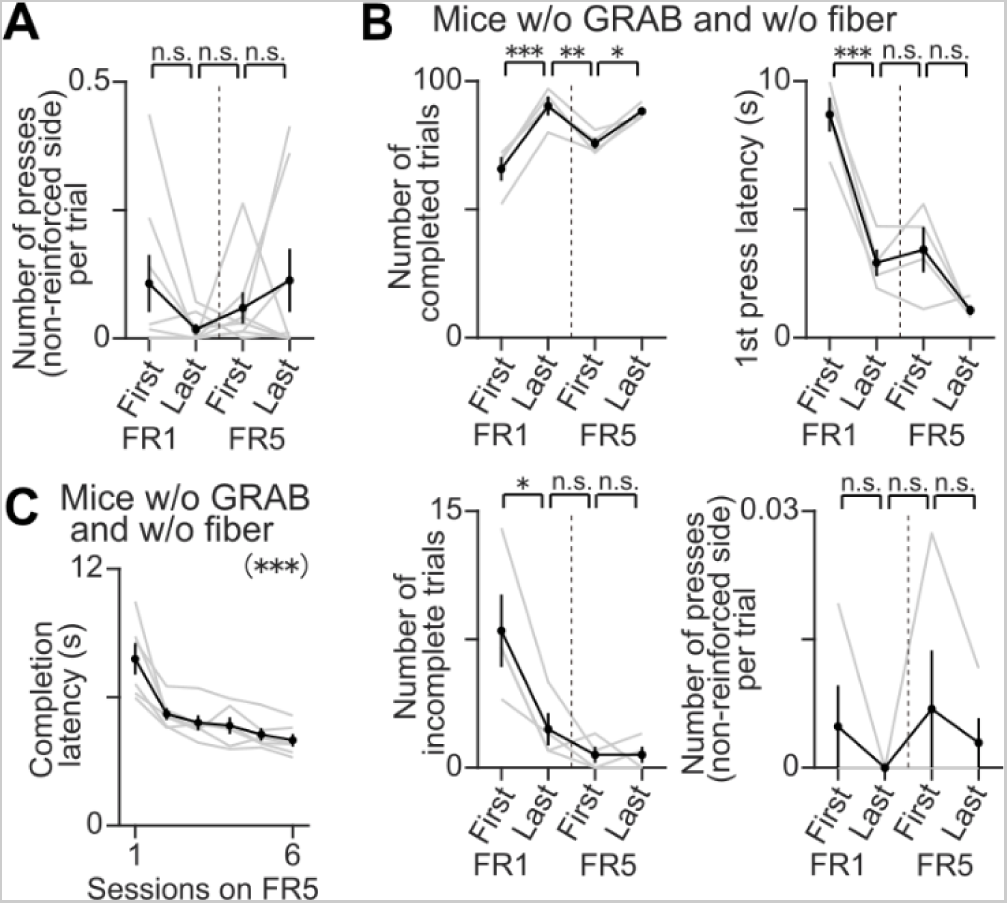
The number of non-reinforced lever presses shows no significant changes in a session-by-session comparison. (A) Session-by-session comparison of the number of lever presses on the other side, which was not reinforced in the task, per trial among four selected sessions: FR1-First, FR1-Last, FR5-First, and FR5-Last. The schedule transition from the FR1 to the FR5 schedule happened between FR1-Last and FR5-First, as indicated by the vertical dashed line (n = 4 sessions with 8 mice each, Tukey-Kramer post-hoc analysis, FR1-First versus FR1-Last, p = 0.43; FR1-Last versus FR5-First, p = 0.89; FR5-First versus FR5-Last, p = 0.79). Gray line, individual mouse. Black line, mean ± SEM. (B) Session-by-session comparison of the behavior of another group of mice without GRAB_DA2m_ expression and without optic fiber implantation, related to Fig. 1D and Extended Data Fig. 1-1A. These mice showed similar characteristic behavior changes as the GRAB_DA2m_-expressing mice in terms of the number of completed trials, first lever press latency, number of incomplete trials, and number of non-reinforced lever presses among the four selected sessions around the schedule transition (vertical dashed line) (n = 4 sessions with 8 mice each, Tukey-Kramer post-hoc analysis, FR1-First versus FR1-Last, p = 2.4 × 10^−4^ [number of completed trials], p = 1.7 × 10^−4^ [first press latency], p = 1.1 × 10^−2^ [number of incomplete trials], p = 0.66 [number of presses on unreinforced side]; FR1-Last versus FR5-First, p = 9.3 × 10^−3^ [number of completed trials], p = 0.91 [first press latency], p = 0.70 [number of incomplete trials], p = 0.39 [number of presses on unreinforced side]; FR5-First versus FR5-Last, p = 2.2 × 10^−2^ [number of completed trials], p = 5.3 × 10^−2^ [first press latency], p > 0.99 [number of incomplete trials], p = 0.78 [number of presses on unreinforced side]). Gray line, individual mouse. Black line, mean ± SEM. * p < 0.05. ** p < 10^−2^. *** p < 10^−3^. (C) Completion latency, which is the duration between the first and last lever presses (Fig. 2B, right), detected behavior adjustment of the group of mice without GRAB_DA2m_ expression and without optic fiber implantation as well as the GRAB_DA2m_-expressing mice during the FR5 sessions (Fig. 1E) (n = 6 sessions with 6 mice each, one-way repeated measures ANOVA, *F*_(5,25)_ = 24.00, p = 8.5 × 10^−9^). Gray line, individual mouse. Black line, mean ± SEM. *** p < 10^−8^.

**Extended Data Figure 2-1.**
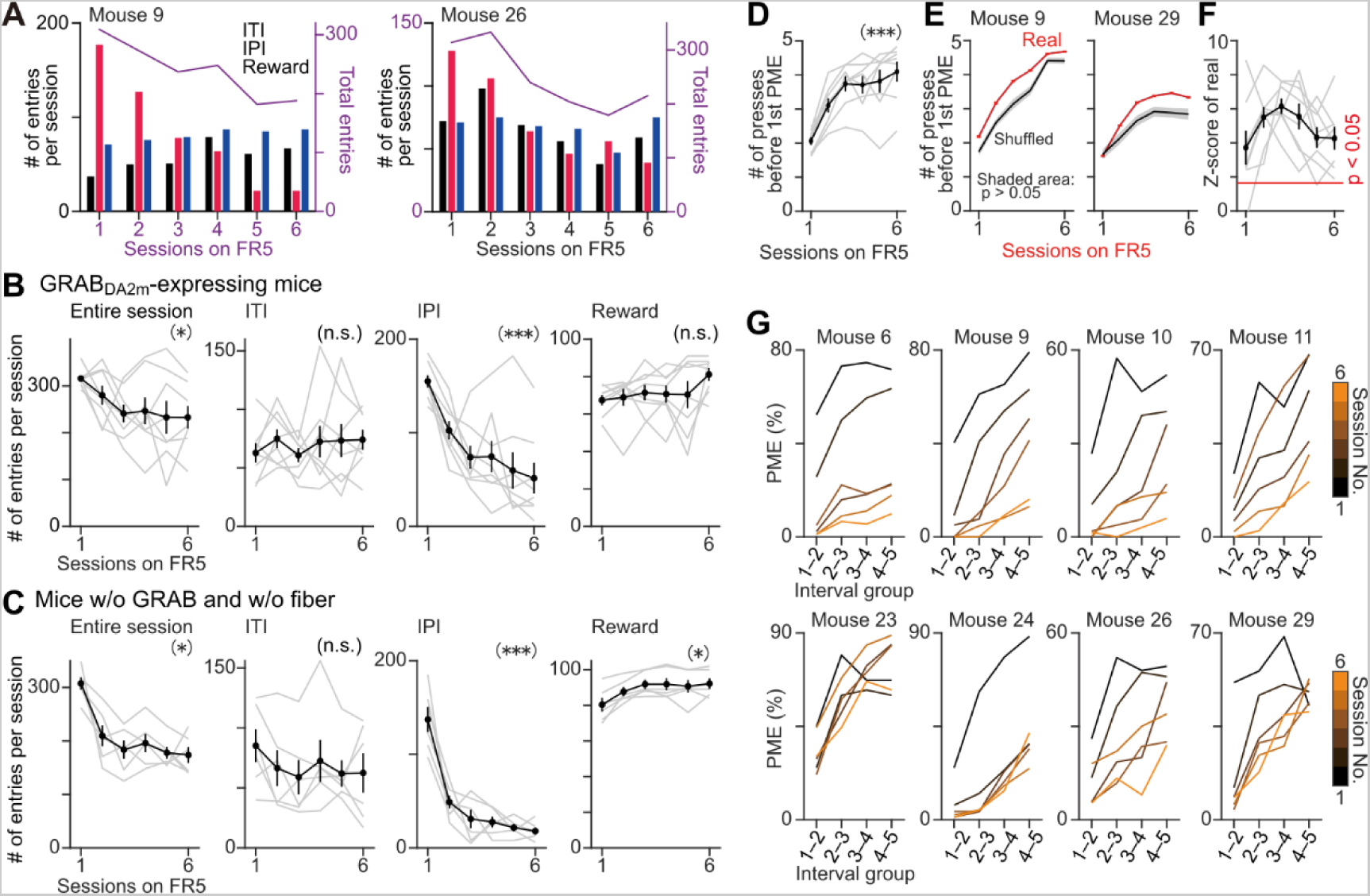
Progressive reduction in the total number of head entries across the FR5 sessions is explained by the reduction in the number of PMEs. (A) Session-by-session comparison of total head entries into the magazine (purple line) and comparisons of the number of head entries during the ITI (black bar), inter-press interval (IPI, red bar), or reward period (blue bar) across the FR5 sessions in two representative mice. Note the gradual decrease in the number of entries during the inter-press interval period corresponding to the change in the total head entries, while there are no such decreases in the number during the other two periods. (B) Group data showing the session-by-session changes in the number of head entries to the magazine across the FR5 sessions in all mice. Leftmost: The number of total head entries progressively reduced. Second from left: The number of head entries in the ITI period stayed relatively constant. Second from right: Only the number of head entries during the inter-press interval period significantly reduced, corresponding to the decrease in the total head entries. Rightmost: The number of head entries at the reward period staying relatively constant (n = 6 sessions with 8 mice each, one-way repeated measures ANOVA, entire session, *F*_(5,35)_ = 3.00, p = 2.4 × 10^−2^; ITI, *F*_(5,35)_ = 0.47, p = 0.80; IPI, *F*_(5,35)_ = 11.98, p = 8.4 × 10^−7^; reward, *F*_(5,35)_ = 1.62, p = 0.18). Gray line, individual session. Black line, mean ± SEM. * p < 0.05. *** p < 10^−6^. (C) Group data showing the session-by-session changes in the number of head entries to the magazine across the FR5 sessions in another group of mice without GRAB_DA2m_ expression and without optic fiber implantation, related to Extended Data Fig. 2-1B. Similar to the GRAB_DA2m_-expressing mice, the number of total head entries to the magazine progressively decreased (leftmost panel), and only the entries during the inter-press interval period (second from the right) accounted for this decrease (n = 6 sessions with 8 mice each, one-way repeated measures ANOVA, entire session, *F*_(5,25)_ = 16.06, p = 4.1 × 10^−7^; ITI, *F*_(5,25)_ = 1.58, p = 0.20; IPI, *F*_(5,25)_ = 29.29, p = 1.1 × 10^−9^; reward, *F*_(5,25)_ = 10.80, p = 1.3 × 10^−5^). Gray line, individual session. Black line, mean ± SEM. * p < 10^−4^. *** p < 10^−8^. (D) The average number of lever presses before the mice made their first PME or ME in a given trial, a behavioral readout reflecting how many presses the mice expected as adequate for a reward, progressively increased across the FR5 sessions in all mice tested (n = 6 sessions with 8 mice each, one-way repeated measures ANOVA, entire session, *F*_(5,35)_ = 29.16, p = 1.5 × 10^−11^). *** p < 10^−10^. (E) The average number of lever presses before their first PME or a ME in a given trial (red line, mean ± SEM) was significantly larger than those with PME group number-shuffled counterparts (black line and grey shaded data, mean ± 1.65 standard deviation [p > 0.05], shuffled 1000 times) in two representative mice, suggesting that the mice recognize the press number or PME group number and organize the timings of their reward-seeking behavior accordingly throughout the FR5 sessions. (F) Group data of the z-scored number of lever presses before a first PME or a ME in a given trial— calculated based on the corresponding shuffled datasets—across the FR5 sessions in all mice. Z-scores of 46 sessions out of 48 sessions corresponded to p < 0.05 (red line). (G) Session-by-session plots of the PME percentage versus the interval group during the FR5 schedule of all mice, related to Fig. 2G (n = 24 data points, 4 interval groups × 6 sessions, two-way factorial ANOVA, interval group number, *F*_(3,23)_ = 17.96, p = 3.2 × 10^−5^ [mouse 6], *F*_(3,23)_ = 18.48, p = 2.7 × 10^−5^ [mouse 9], *F*_(3,23)_ = 8.49, p = 1.6 × 10^−3^ [mouse 10], *F*_(3,23)_ = 28.83, p = 1.8 × 10^−6^ [mouse 11], *F*_(3,23)_ = 28.62, p = 1.9 × 10^−6^ [mouse 23], *F*_(3,23)_ = 25.34, p = 4.0 × 10^−6^ [mouse 24], *F*_(3,23)_ = 14.87, p = 9.2 × 10^−5^ [mouse 26], *F*_(3,23)_ = 11.98, p = 2.9 × 10^−4^ [mouse 29]; session number, *F*_(5,23)_ = 96.90, p = 7.1 × 10^−11^ [mouse 6], *F*_(5,23)_ = 25.64, p = 7.5 × 10^−7^ [mouse 9], _(5,23)_ = 19.56, p = 4.3 × 10^−6^ [mouse 10], *F*_(5,23)_ = 22.88, p = 1.6 × 10^−6^ [mouse 11], *F*_(5,23)_ = 3.18, p = 3.8 × 10^−2^ [mouse 23], *F*_(5,23)_ = 24.76, p = 9.5 × 10^−7^ [mouse 24], *F*_(5,23)_ = 12.95, p = 5.3 × 10^−5^ [mouse 26], *F*_(5,23)_ = 4.87, p = 7.6 × 10^−3^ [mouse 29]).

**Extended Data Figure 2-2.**
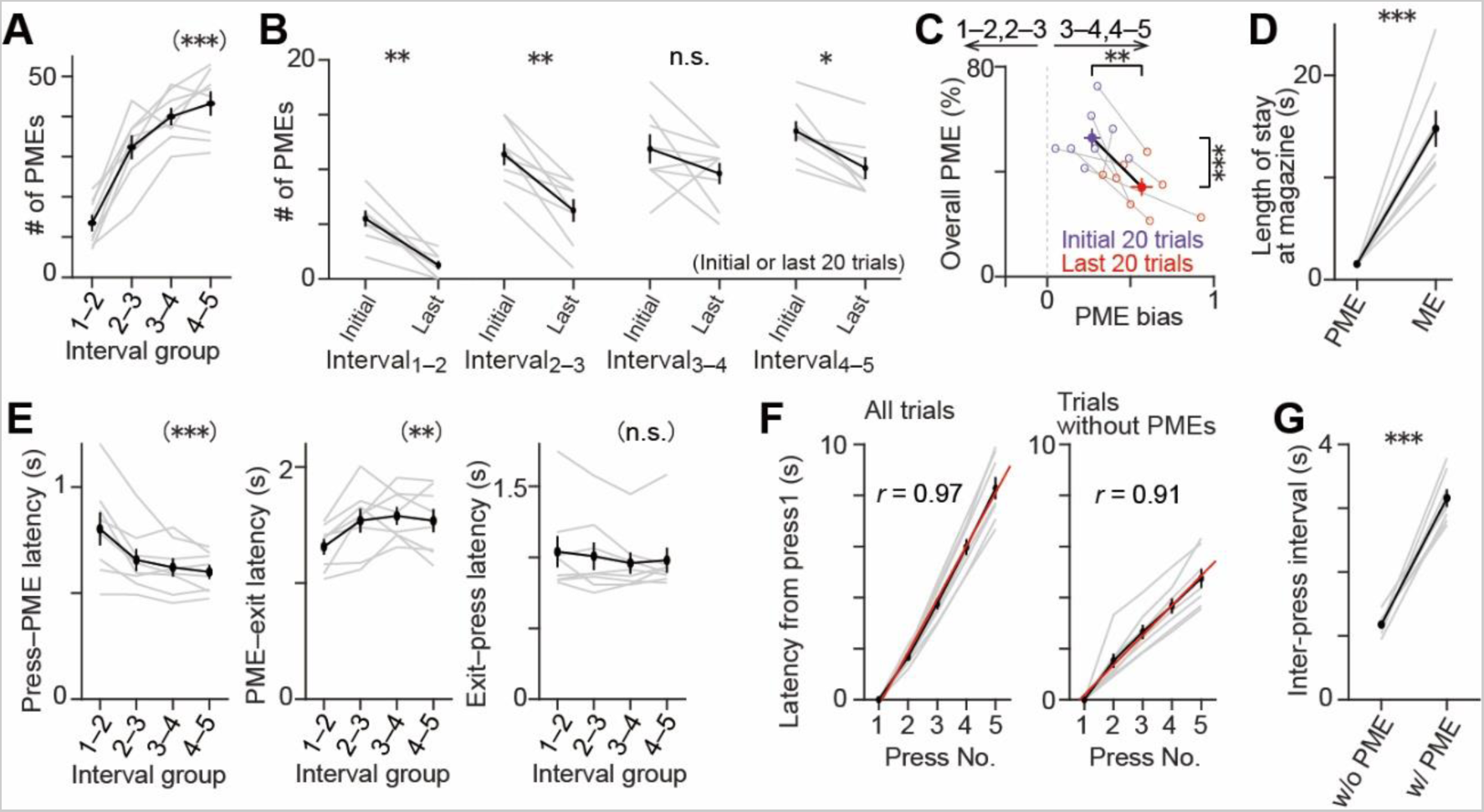
Analysis of the single FR5 session from each mouse selected for further behavioral or fiber photometry analyses. (A) Group data showing that the number of PMEs increased with the increase in the interval group number in the selected FR5 sessions in all mice (n = 4 interval groups with 8 sessions each from 8 mice, number of PMEs = 13.5 ± 2.0 [Interval_2–3_], 32.4 ± 2.9 [Interval_2–3_], 40.0 ± 2.2 [Interval_3–4_], 43.3 ± 3.0 [Interval_4–5_], one-way repeated measures ANOVA, *F*_(3,21)_ = 71.25, p = 3.6 × 10^−11^). The minimum number of events at a PME group was 7. Gray line, individual session. Black line, mean ± SEM. *** p < 10^−10^. (B) Group data of a within-session comparison of the number of events in each PME group between the initial and final 20 trials in the selected FR5 sessions showing a consistent decrease in all mice (n = 8 trial groups from 8 mice each, number of PMEs = 5.5 ± 0.7 [Interval_1–2_, initial trials], 1.3 ± 0.4 [Interval_1–2_, last trials], 11.4 ± 1.0 [Interval_2–3_, initial trials], 6.3 ± 1.0 [Interval_2–3_, last trials], 11.9 ± 1.3 [Interval_3–4_, initial trials], 9.6 ± 1.0 [Interval_3–4_, last trials], 13.5 ± 0.9 [Interval_4–5_, initial trials], 10.1 ± 1.0 [Interval_4–5_, last trials], paired t-test with Bonferroni correction, Interval_1–2_, *t*_(7)_ = 5.86, p = 2.5 × 10^−3^; Interval_2–3_, *t*_(7)_ = 5.09, p = 5.7 × 10^−3^; Interval_3–4_, *t*_(7)_ = 1.46, p = 0.75; Interval_4–5_, *t*_(7)_ = 4.22, p = 0.016). Filled circle, mean ± SEM within a group of trials. Gray line, individual mouse. Black line, mean ± SEM. * p < 0.05.** p < 10^−2^. (C) Group data plot showing that the increase in the PME bias and decrease in the overall PME percentage not only happened across sessions (Fig. 2H) but also within the selected FR5 session (n = 2 trial groups with 8 mice each, PME bias, 0.27 ± 0.05 [initial trials], 0.57 ± 0.07 [last trials], paired t-test, *t*_(7)_ = –3.65, p = 8.2 × 10^−3^; overall PME percentage, 52.8 ± 3.6% [initial trials], 34.1 ± 3.4% [last trials], paired t-test, *t*_(7)_ = –5.58, p = 8.3 × 10^−4^). Gray line, individual mouse. Black line, mean of all mice. ** p < 10^−2^.*** p < 10^−3^. (D) The mice stayed longer at the food magazine after an ME than after a PME (n = 2 groups with 8 mice each, PME, 1.53 ± 0.08 s; ME, 14.79 ± 1.76 s; paired t-test, *t*_(7)_ = – 7.37, p = 1.5 × 10^−4^). Gray line, individual mouse. Black line, mean ± SEM. *** p < 10^−3^. (E) Inter-press intervals with a PME were divided into three parts (press–PME, PME–exit, and exit–press periods). Left: Decreasing trend for the press-to-PME latency with an increasing PME group number (n = 4 interval groups with 8 sessions each from 8 mice, one-way repeated measures ANOVA, *F*_(3,21)_ = 9.82, p = 3.0 × 10^−4^). Center: Significant differences in the length of stay at the magazine following PMEs in the inter-press interval groups, suggesting that it took more time for the mice to realize the same outcome of reward omission with a greater degree of reward expectations (n = 4 interval groups with 8 sessions each from 8 mice, one-way repeated measures ANOVA, *F*_(3,21)_ = 5.28, p = 7.2 × 10^−3^). Right: No significant differences in the latency of a subsequent press following the preceding magazine exit moment (n = 4 interval groups with 8 sessions each from 8 mice, one-way repeated measures ANOVA, *F*_(3,21)_ = 1.56, p = 0.23). Gray line, individual session. Black line, mean ± SEM. ** p < 10^−2^.*** p < 10^−3^. (F) The lever press number was positively correlated with the elapsed time, since the first press in a given trial (regardless of the trials with PMEs) is included (left panel) or not (right panel) (n = 5 press numbers with 8 sessions each from 8 mice, Pearson correlation coefficient, all trials, *r* = 0.97, p = 2.0 × 10^−25^; trials without PMEs, *r* = 0.91, p = 2.7 × 10^−16^). (G) Mice’s making a PME significantly elongated the duration of the corresponding inter-press interval (n = 2 groups with 8 sessions each from 8 mice, without PME, 1.18 ± 0.05 s; with PME, 3.16 ± 0.13 s; paired t-test, *t*_(7)_ = –12.53, p = 4.8 × 10^−6^).

**Extended Data Figure 3-1.**
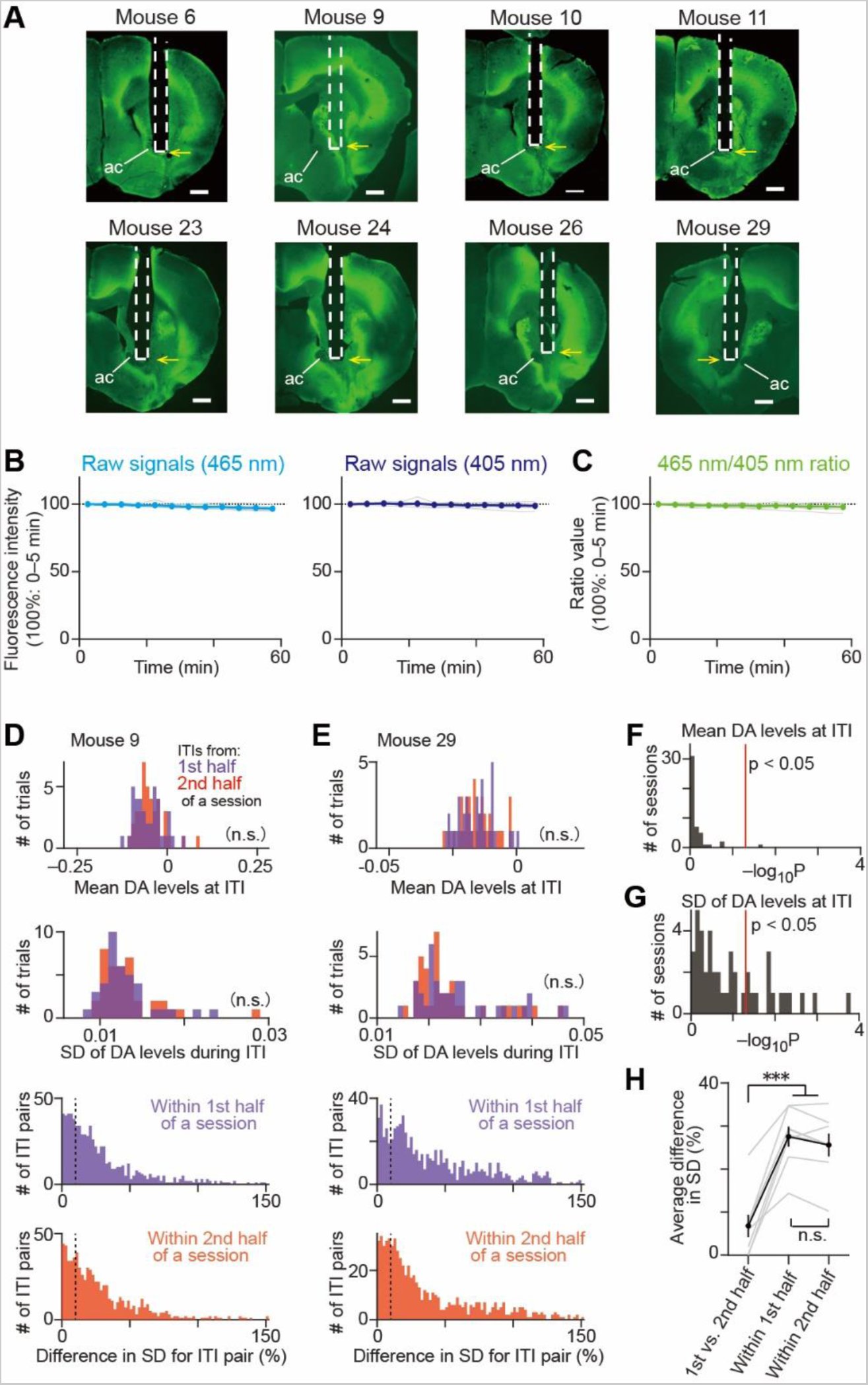
Stable fluorescence intensity for an hour enabled stable *in vivo* DA measurements from freely behaving mice. (A) Coronal brain section showing the location of the optic fiber and the GRAB_DA2m_ expression in the VS of each mice tested. Dashed line, fiber track. ac: anterior commissure. Yellow arrow, fiber tip. (B) Fluorescence intensity variations by 405 nm (left) or 465 nm (right) excitation over one hour. We observed no significant fluorescence intensity reduction due to photobleaching in any signals at any periods (n = twelve 5-min bins with 6 sessions each from 3 mice, 2 sessions from each mice, one-sample t-test with Bonferroni correction, 405 nm excitation, fluorescence intensity > 98.8 ± 1.1%, all *t*_(5)_ > –1.17, all p > 0.99; 465 nm excitation, fluorescence intensity > 96.7 ± 0.8%, all *t*_(5)_ > –4.13, all p > 0.11). Note that the mice were individually placed in an operant chamber but not subjected to a task. Horizontal dotted line, 100% fluorescence intensity (0–5 min). Gray line, individual session. Thicker line with filled circles, mean ± SEM. (C) Same as (B) but for the 465 nm/405 nm ratio. We observed no significant reduction in ratio from 100% at any periods (n = twelve 5-min bins with 6 sessions each from 3 mice, 2 sessions from each mice, one-sample t-test with Bonferroni correction, ratio > 98.0 ± 1.2%, all *t*_(5)_ > –1.75, all p > 0.99). (D) Investigation of the progressive reduction in the DA signal fluctuation ranges between the first and second halves of the trials of a session in a representative mouse. Top: Comparison of the distributions of the average DA signal levels in the 465 nm/405 nm ratio (high-pass filtered at 0.0167 Hz) in each ITI period between the first and the second halves of a session, showing no significant difference (n = 38 trials each, Kolmogorov-Smirnov test, *D_max_* = 0.11, p = 0.64). Second from top: Same as the top panel but for the comparison of the distributions of the standard deviation, the results of which exclude the possibility of dynamic changes in the standard deviation of the DA release (*D_max_* = 0.11, p = 0.64). Bottom two panels: Differences in the SD within each half of a session, suggesting large within-section variations in the SD (n = 703 trial pairs from 38 trials, first half, 71.4% of pairs (502 trials) with more than a 10% difference; second half, 72.8% of pairs (512 trials) with more than a. 10% difference). Vertical dashed line, 10% difference. (E) Same as ***Extended Data Fig. 2-2D*** but for the investigation of the progressive reduction in the DA signal fluctuation ranges in another representative mouse. Top: Comparison of the distributions of the average DA signal levels (n = 38 and 37 trials for the first and second halves, respectively, Kolmogorov-Smirnov test, *D_max_* = 0.05, p = 0.90). Second from top: Same as the top panel but for the comparison of the distributions of the SD (*D_max_* = 0.19, p = 0.22). Bottom two panels: Differences in the SD within each half of a session (n = 703 trial pairs from 38 trials, first half, 80.8% of pairs (568 trials) with more than 10% difference; second half, 75.8% of pairs (505 trials) with more than 10% difference). Vertical dashed line, 110% difference. (F) In the majority of the sessions, we did not find significant reductions in the mean DA levels at ITI periods between the first and second halves of a session (Kolmogorov-Smirnov test, p > 0.05 in 47 out of 48 sessions from 8 mice). Red line: p = 0.05. (G) In the majority of the sessions, we did not find significant reductions in the SD of the DA levels at ITI periods between the first and second halves of a session (Kolmogorov-Smirnov test, p > 0.05 in 30 out of 48 sessions from 8 mice). Red line: p = 0.05. (H) We found significantly large differences in the SDs of the DA levels in the ITIs within the first and second halves of the sessions compared to the differences caused by a progressive change in the SDs between the first and second sessions. Since the normalization of the DA signals for the z-score calculation using SDs varying among the trials rather than across the sections may impair the comparison of the DA response amplitude throughout a session, these results encouraged us to utilize a uniform normalization factor for the z-score calculation for each session (n = 3 groups with 8 mice each, between halves (first versus second halves), 6.8 ± 2.6%, within first half, 27.5 ± 2.3%, within second half, 25.6 ± 2.6%, Tukey-Kramer post-hoc analysis, between halves vs within first half, p = 1.6 × 10^−6^; between halves vs within second half, p = 5.1 × 10^−6^; within 1st half vs within 2nd half, p = 0.71). Gray line, individual mouse. Black line, mean ± SEM.

**Extended Data Figure 3-2.**
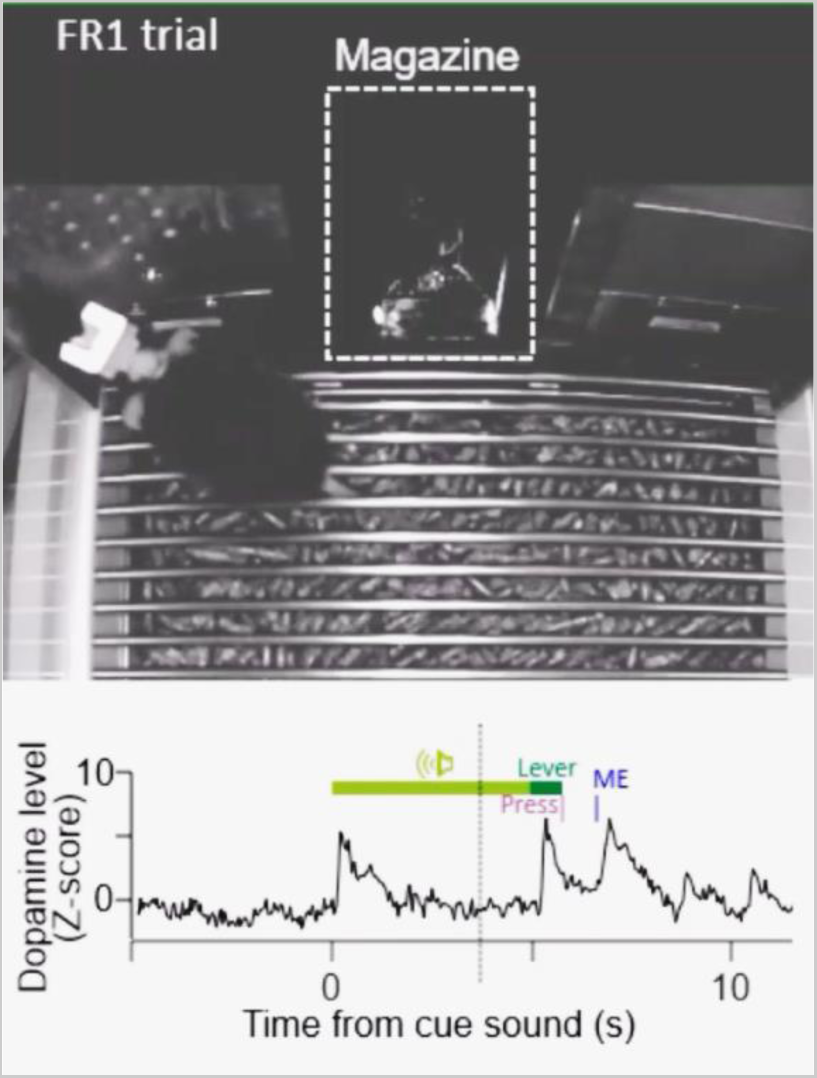
Mouse behavior and extracellular dopamine fluctuations during the FR1 task. (Top) Representative behavior of a mouse engaging in the FR1 task. After the 5-s noise sound cue, reinforced (left) and non-reinforced (right) levers were presented. The mouse was rewarded at the magazine (white dashed line) after pressing the reinforced lever once per trial. (Bottom) Simultaneous measurement of extracellular DA fluctuations in the right ventral striatum. The vertical dotted line moves according to the camera frame timing. A phasic DA increase was observed upon the onset of a sound cue (light green bar), lever extension (dark green bar), or ME (blue tick).

**Extended Data Figure 4-1.**
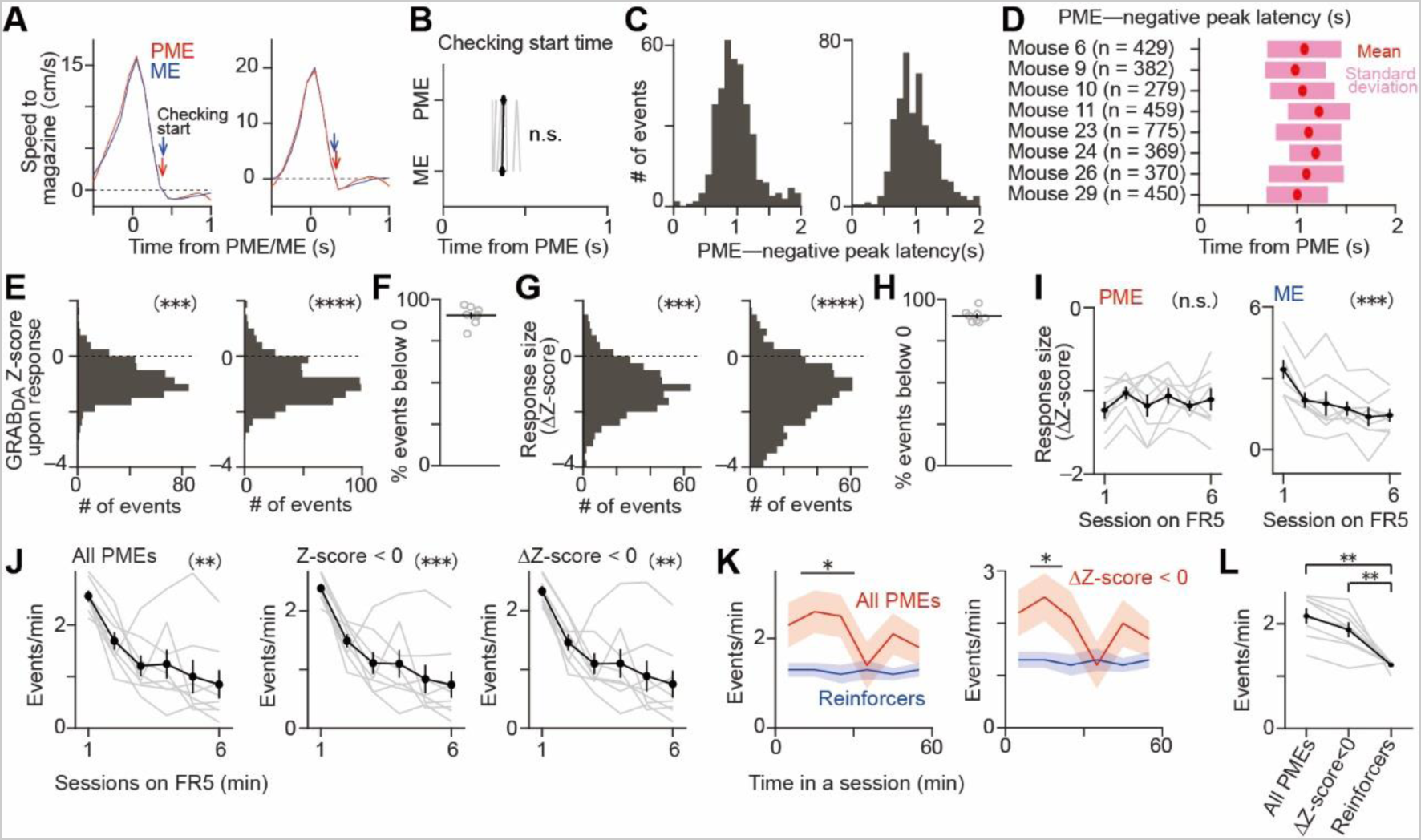
PMEs induce DA dips. (A) Average changes in the head speed toward the magazine around the PME or ME in two representative mice. A positive value corresponds to mice’s approaching the magazine. We defined the putative onset of mice’s magazine checking behavior as the time when the head speed became equal to or less than 0 cm/s following the entry. Arrowhead, putative starting timing of magazine checking. Red line and shade, PME. Blue line and shade, ME. Data shown as mean ± SEM. Left panel: Mouse 10, n = 397 PMEs and 403 MEs from 6 FR5 sessions. Right panel: Mouse 23, n = 861 PMEs and 360 MEs from 6 sessions. (B) Group data showing the putative checking start time after a PME/ME in all mice. According to these results, we set the baseline period for the calculation of the DA responses upon a PME/ME as 0–0.2 s (n = 2 data groups with 8 mice each, PME, 0.37 ± 0.02 s, ME, 0.36 ± 0.02, paired t-test, *t*_(7)_ = 1.30, p = 0.24). Gray line, individual mouse. Black line, mean ± SEM. (C) Temporal distributions of the negative peaks of DA responses following PMEs in two representative mice. The responses with a large amplitude (Δz-score < –2.33) at the peak were selected and analyzed. Left panel: Mouse 9, n = 382 events from 6 sessions, 0.98 ± 0.31 s (mean ± standard deviation). Right panel: Mouse 29, n = 450 events from 6 sessions, 1.00 ± 0.31 s (mean ± standard deviation). (D) The average latency and standard deviation of the negative peak in the DA responses following PMEs in all mice recorded. According to these results, we set the period for the calculation of a DA response upon a PME as 0.5–1.5 s. Average latency, 1.09 ± 0.03 s (n = 8 mice, mean ± SEM). Filled circle, mean. Shade, standard deviation. (E) Distributions of the z-scores in the DA response period (0.5–1.5 s following a PME, refer to the ***Extended Data Fig. 4-1D***) in two representative mice. Left panel: Mouse 9, n = 490 events from 6 sessions, z-score = –0.85 ± 0.03. We found that 90.8% of the responses (445 events) showed z-scores below zero (Wilcoxon signed rank test, signed rank = 3957, Z = –17.92, p = 8.5 × 10^−72^). Right panel: Mouse 29, n = 648 events from 6 sessions, z-score = –0.95 ± 0.03. 89.8% (582 events) showed z-scores below zero (signed rank = 9596, Z = –20.04, p = 2.4 × 10^−89^). *** p < 10^−71^. **** p < 10^−88^. (F) Group data showing the percentage of DA responses with z-scores below zero in all mice recorded. 90.3 ± 2.0% (n = 8 mice). (G) Same as ***Extended Data Fig. 4-1E*** but for distributions of the DA response amplitude (Δz-score) in two representative mice. Left panel: Mouse 9, n = 490 events from 6 sessions, z-score = –1.15 ± 0.04. 90.2% of the responses (442 events) showed negative deflections (Δz-score < 0) (Wilcoxon signed rank test, signed rank = 4646, Z = –17.70, p = 4.3 × 10^−70^). Right panel: Mouse 29, n = 648 events from 6 sessions, z-score = –1.37 ± 0.05. 88.7% of the responses (575 events) showed negative deflections (signed rank = 8992, Z = –20.17, p = 1.9 × 10^−90^). *** p < 10^−69^. **** p < 10^−89^. (H) Group data showing the percentage of DA responses with negative deflections (Δz-score < 0) in all mice recorded. 89.94 ± 1.35% (n = 8 mice). (I) No significant changes in the response amplitude upon a PME (left panel) but upon a ME (right panel) across the FR5 sessions in all mice recorded (n = 6 sessions with 8 mice each, one-way repeated measures ANOVA, PME, *F*_(5,35)_ = 1.68, p = 0.17; ME, *F*_(5,35)_ = 25.42, p = 9.7 × 10^− 11^), suggesting that a DA response upon a PME in any FR5 session has the potential to serve as, in the RL theory, a teaching signal with a similar weight throughout the FR5 sessions. Gray line, individual mouse. Black line, mean ± SEM. *** p < 10^−10^. (J) Group data showing progressive reductions in the frequency of all PMEs (left), PMEs followed by a DA response with the z-score < 0 (center), and those followed by a negative deflection (Δz-score < 0) across the FR5 sessions in all mice (n = 6 sessions with 8 mice each, one-way repeated measures ANOVA, all PMEs, *F*_(5,35)_ = 11.82, p = 9.7 × 10^−7^; PMEs with DA responses with negative z-scores, *F*_(5,35)_ = 17.98, p = 8.2 × 10^−9^; PMEs with DA responses with negative deflections, *F*_(5,35)_ = 13.18, p = 3.0 × 10^−7^). Gray line, individual mouse. Black line, mean ± SEM. ** p < 10^−6^. *** p < 10^−8^. (K) Within-session changes in the frequency of all PMEs (left, red line) and those followed by a negative deflection in the DA signal (right, red line) plotted together with the frequency of task accomplishments (blue line) in the selected FR5 session of a representative mouse. Bar with an asterisk, p < 0.05 (paired t-test with Bonferroni correction). (L) Group data showing that the frequency of all PMEs and PMEs with DA responses with negative deflections is higher than that of the mice’s obtaining reinforcers in the selected FR5 sessions of all mice (n = 3 data groups with 8 mice each, number of events per minute = 2.2 ± 0.2 [all PMEs], 1.9 ± 0.1 [responses with Δz -score < 0], 1.2 ± 0.0 [reinforcers], paired t-test with Bonferroni correction, all PMEs versus reinforcers, *t*_(7)_ = 7.72, p = 1.4 × 10^−3^, responses with Δz -score < 0 versus reinforcers, *t*_(7)_ = 4.04, p = 9.8 × 10^−3^). Gray line, individual mouse. Black line, mean ± SEM. ** p < 10^−2^.

**Extended Data Figure 4-2.**
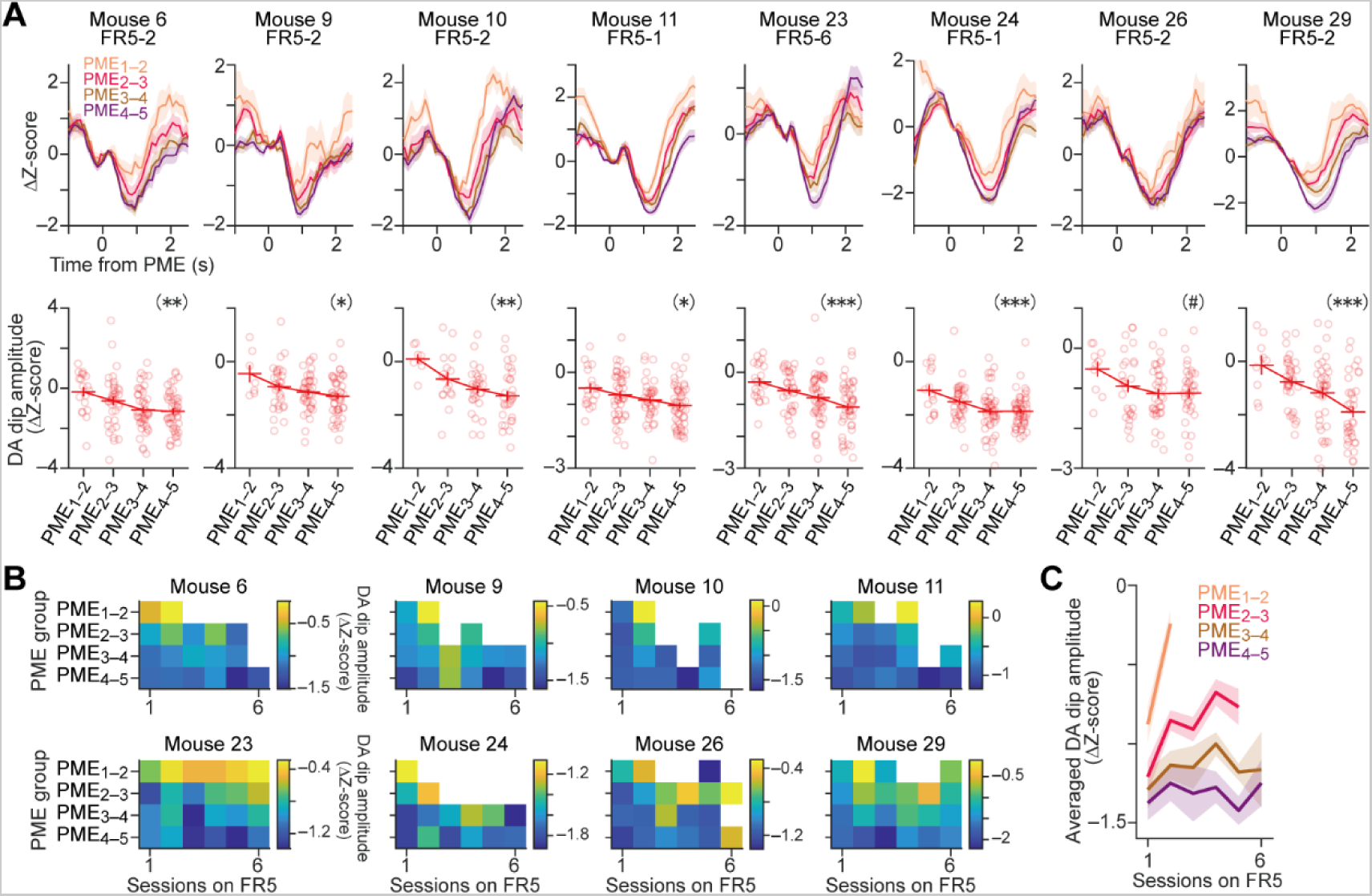
Plots of DA dip graduality upon PMEs in all mice recorded. (A) GRAB_DA2m_ traces (top row) and corresponding statistics (bottom row) of response amplitude for each PME group in the selected FR5 sessions of all the mice recorded (related to Fig. 4F). The magnitude of the DA response following a PME consistently increased with the increasing PME group number (one-way ANOVA, mouse 6, *F*_(3,146)_ = 4.61, p = 4.1 × 10^−3^; mouse 9, *F*_(3,126)_ = 3.28, p = 2.3 × 10^−2^; mouse 10, *F*_(3,84)_ = 5.66, p = 1.4 × 10^−3^; mouse 11, *F*_(3,150)_ = 3.87, p = 1.1 × 10^−2^; mouse 23, *F*_(3,146)_ = 6.79, p = 2.6 × 10^−4^; mouse 24, *F*_(3,151)_ = 6.26, p = 5.0 × 10^−4^; mouse 26, *F*_(3,105)_ = 2.28, p = 0.084; mouse 29, *F*_(3,117)_ = 8.27, p = 5.0 × 10^−5^). # p < 0.1. * p < 0.05. ** p < 10^−2^. *** p < 10^−3^. (B) Color plot for each mouse showing the progressive change in the DA dip amplitude for each PME group across the FR5 sessions. Grids with less than seven DA dip events (area in white) are excluded from the plot. (C) Group data comparison of the DA dip amplitude in each PME group across the FR5 sessions, related to ***Extended Data Figure 4-2B***. n = 4–8 mice each at each data point. Line and shade: mean ± SEM.

**Extended Data Figure 4-3.**
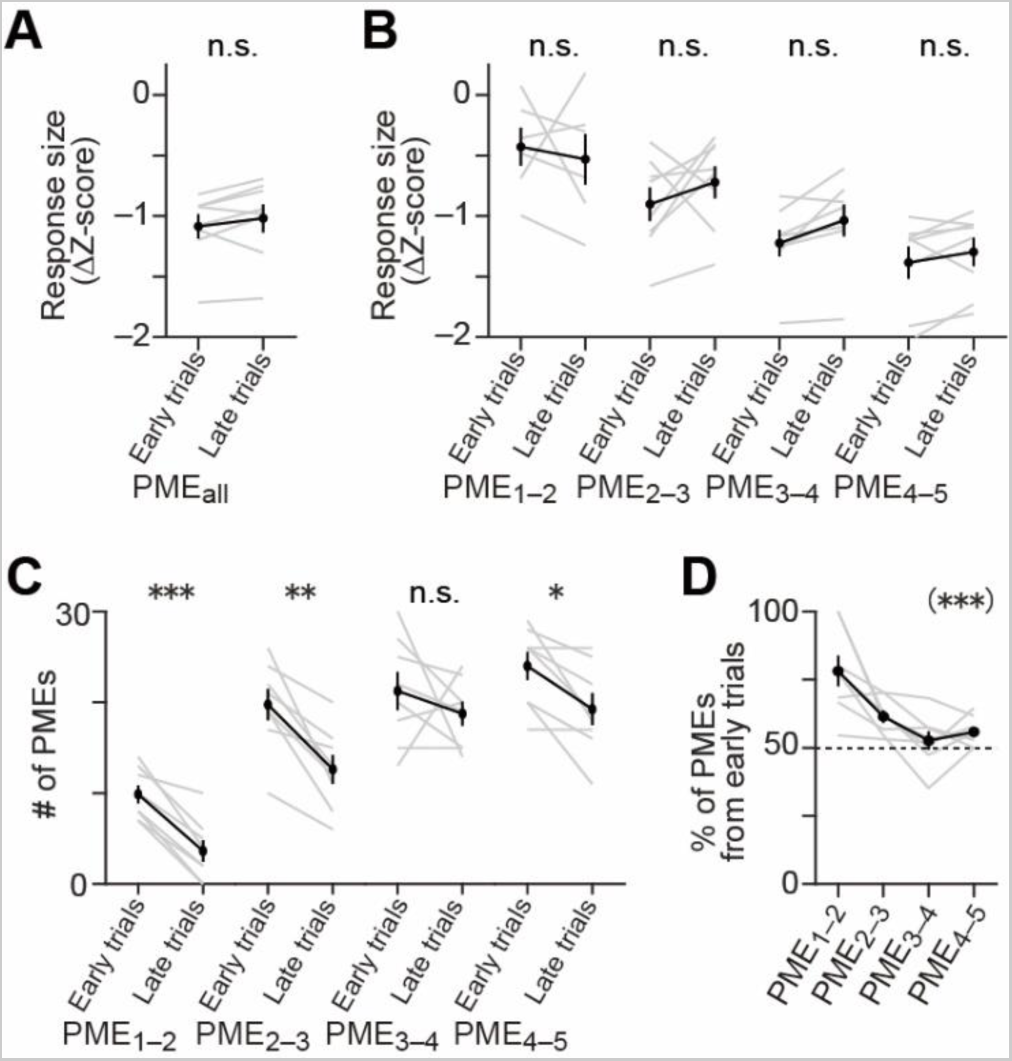
No significant within-session progressive changes in the DA dip amplitude upon PMEs. (A–D) Dopaminergic and behavioral analyses excluding the possibility that within-session progressive reductions in the DA responses upon PMEs within a session may explain the variability in the DA dip amplitude. (A) Group data showing that the DA response (dip) amplitude was not consistently changed between the early trials (first half) and late trials (second half) in the selected FR5 sessions of all mice. The result contradicted our hypothesis that the amplitude would progressively decrease as the mice became familiar with the task (n = 8 sessions each from 8 mice, response amplitude (Δz-score) = –1.09 ± 0.10 [early trials], –1.02 ± 0.12 [late trials]; paired t-test, *t*_(7)_ = –1.45, p = 0.19). Gray line, individual mouse. Black line, mean ± SEM. (B) Related to ***Extended Data Fig. 4-3A*** but for the absence of significant progressive reductions in the DA response amplitude for each PME group, suggesting that such progressive reductions cannot explain the variability in the DA dip amplitude among the PME groups (n = 6– 8 sessions each from 6–8 mice, response amplitude (Δz-score) = –0.43 ± 0.16 [PME_1–2_, early trials], –0.53 ± 0.21 [PME_1–2_, late trials], –0.90 ± 0.14 [PME_2–3_, early trials], –0.72 ± 0.13 [PME_2–3_, late trials], –1.22 ± 0.11 [PME_3–4_, early trials], –1.04 ± 0.13 [PME_3–4_, late trials], –1.38 ± 0.13 [PME_4–5_, early trials], –1.3 ± 0.12 [PME_4–5_, late trials], paired t-test with Bonferroni correction, PME_1–2_, *t*_(5)_ = 0.42, p > 0.99; PME_2–3_, *t*_(7)_ = –1.08, p > 0.99; PME_3–4_, *t*_(7)_ = –2.94, p = 0.087; PME_4–5_, *t*_(7)_ = –1.32, p = 0.91). Gray line, individual mouse. Black line, mean ± SEM. (C) Group data showing that the number of PMEs significantly and consistently decreased within a session for most PME groups in all mice (n = 8 sessions each from 8 mice, number of PMEs = 9.9 ± 1.0 [PME_1–2_, early trials], 3.6 ± 1.2 [PME_1–2_, late trials], 19.8 ± 1.7 [PME_2–3_, early trials], 12.6 ± 1.6 [PME_2–3_, late trials], 21.3 ± 2.1 [PME_3–4_, early trials], 18.8 ± 1.3 [PME_3–4_, late trials], 24.0 ± 1.6 [PME_4–5_, early trials], 19.3 ± 1.8 [PME_4–5_, late trials], paired t-test with Bonferroni correction, PME_1–2_, *t*_(7)_ = 7.44, p = 5.8 × 10^−4^; PME_2–3_, *t*_(7)_ = 4.66, p = 9.3 × 10^−3^; PME_3–4_, *t*_(7)_ = 0.92, p > 0.99; PME_4–5_, *t*_(7)_ = 3.37, p > 0.048). Gray line, individual mouse. Black line, mean ± SEM. (D) Comparison of the proportions of the PMEs occurred in early trials across the PME groups. Data for the smaller PME group numbers consisted of relatively more events occurring in the early trials, which is against our hypothesis that smaller PME group numbers, inducing smaller DA responses, would be more likely in later trials, the part hypothesized to induce smaller DA dips (n = 8 sessions each from 8 mice, percent of PMEs from early trials, 78.2 ± 5.6% [PME_1–2_], 61.5 ± 2.3% [PME_2–3_], 52.6 ± 3.4% [PME_3–4_], 55.8 ± 1.9% [PME_4–5_], one-way repeated-measures ANOVA, *F*_(3,21)_ = 11.46, p = 1.2 × 10^−4^). Gray line, individual mouse. Black line, mean ± SEM.

**Extended Data Figure 4-4.**
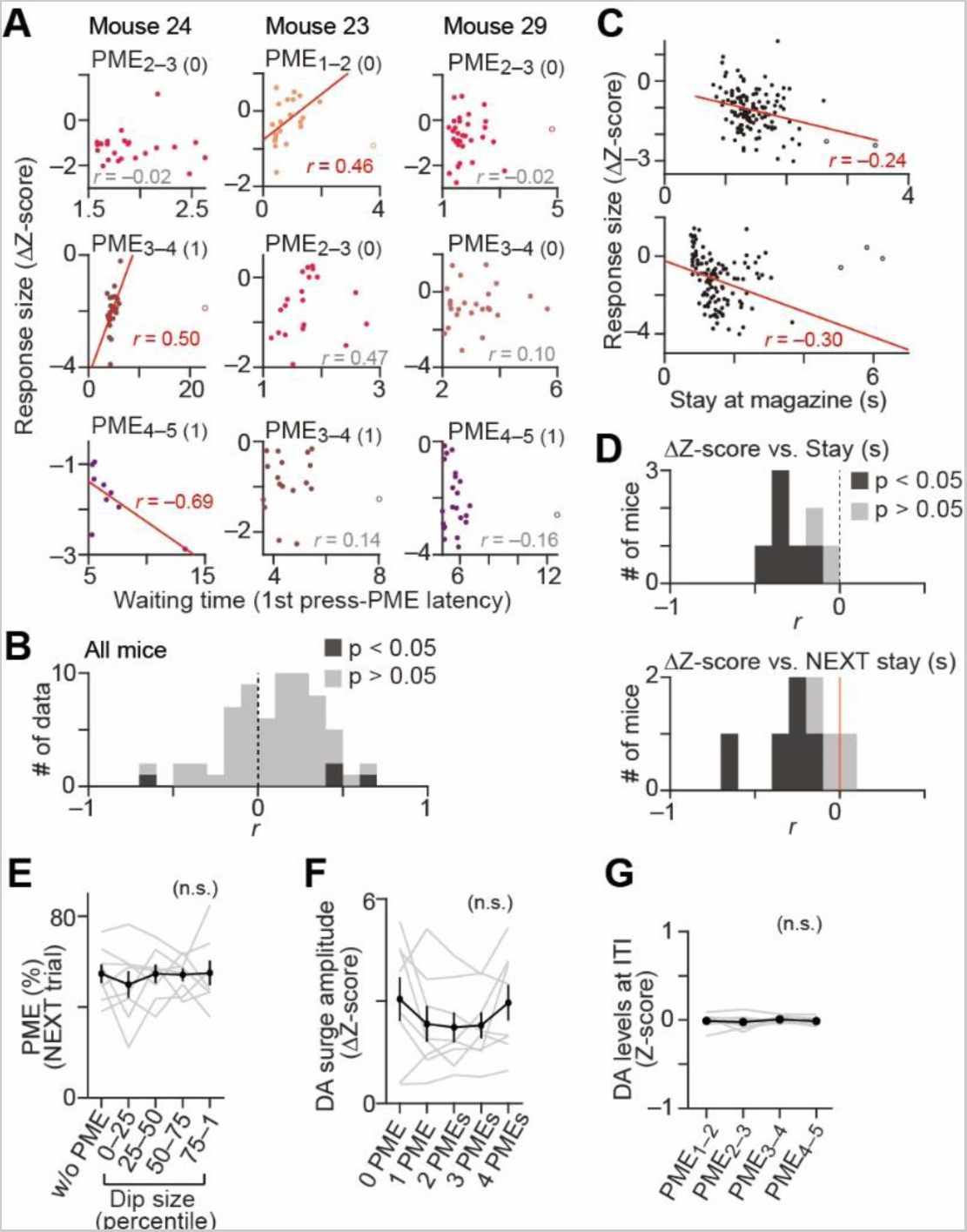
DA dip amplitude graduality upon PMEs is not explained by the elapsed time since mice’s task engagement. (A) Relationship between the latency of a PME from the task engagement (a first press in a given trial) in three representative mice. To exclude the possibility that the DA dip amplitude is correlated with time rather than the number of lever presses, we classified the PME data based on the PME groups (PME_1–2_/PME_2–3_/PME_3–4_/PME_4–5_) and the number of previous PMEs in that trial (number enclosed in parentheses). Three representative data groups from each mouse are shown. Data points outside the mean ± 3 STD were excluded from the correlation analysis (open circles). The Pearson correlation coefficient is shown in each panel. Panel with red line, significant positive or negative correlations. Panel without a line, no significant correlation. (B) Group data related to ***Extended Data Fig. 4-4A*** showing the Pearson correlation coefficients (65 data groups out of 80 groups, with 7 or more samples each) for all data groups of all mice tested. No data groups except one showed a significant negative correlation between the DA dip size and the time elapsed, which excludes the possibility that the waiting time explains the variability in the DA dip amplitude across the PME groups. Shade in dark gray, data group with p < 0.05. Shade in light gray, data group with p > 0.05. (C) Representative data from two mice showing that a longer stay at the magazine upon a PME coincided with a larger (more negative) DA response magnitude (Pearson correlation coefficient, Mouse 9, n = 127 events, *r* = –0.24, p = 8.2 × 10^−3^; Mouse 29, n = 118 events, *r* = –0.30, p = 1.3 × 10^−3^). Data outside the mean ± 3 STD were excluded from the correlation analysis (filled circles). (D) Top: Group data related to ***Extended Data Fig. 4-4C*** showing the Pearson correlation coefficients (8 sessions from 8 mice). Six out of eight mice showed significant negative correlations (p < 0.05, shaded in dark gray), and all mice showed negative correlation coefficients, excluding the possibility that PMEs provided aversive experiences so that the mice quit the magazine entry earlier if the DA responses were more negative. Bottom: Group data same as the top panel but showing the Pearson correlation coefficients between the DA response amplitude upon a PME and the duration of stay at the magazine upon a *subsequent* PME in the same trial. Five out of eight mice showed significant negative correlations (p < 0.05, shaded in dark gray), and seven mice showed negative correlation coefficients, indicating that the duration of stay at the magazine following a bigger DA dip tended to be longer as well. (E) Relationship between the DA dip amplitude upon a PME in a trial and the PME percentage in the next trial. The bigger DA dips did not decrease the PME percentage with the corresponding press number, suggesting that the effects of DA dips on the subsequent behavior cannot be explained by simple framing based on classical reinforcement learning theory (n = 5 data groups with 8 sessions each from 8 mice, one-way repeated-measures ANOVA, *F*_(4,28)_ = 0.39, p = 0.82). The plotted results correspond to the average values among the inter-press interval groups per dip size percentile in the selected FR5 session. Gray line, individual mouse. Black line, mean ± SEM. (F) The DA surge amplitude was not sensitive to how efficiently the mice completed an FR5 trial, suggesting that behavior adjustment during the FR5 sessions was not realized simply via the refinement of performance by reinforcing a good performance with fewer PMEs (n = 5 data groups with 8 sessions each from 8 mice, one-way repeated-measures ANOVA, *F*_(4,28)_ = 1.60, p = 0.20). Gray line, individual mouse. Black line, mean ± SEM. (G) DA levels during the ITI periods (–15 to 0 s to sound cue onsets) did not show significant variances among PME group numbers (n = 8 sessions each from 8 mice, one-way repeated-measures ANOVA, *F*_(3,21)_ = 0.4, p = 0.76). Gray line, individual mouse. Black line, mean ± SEM.

**Extended Data Figure 4-5.**
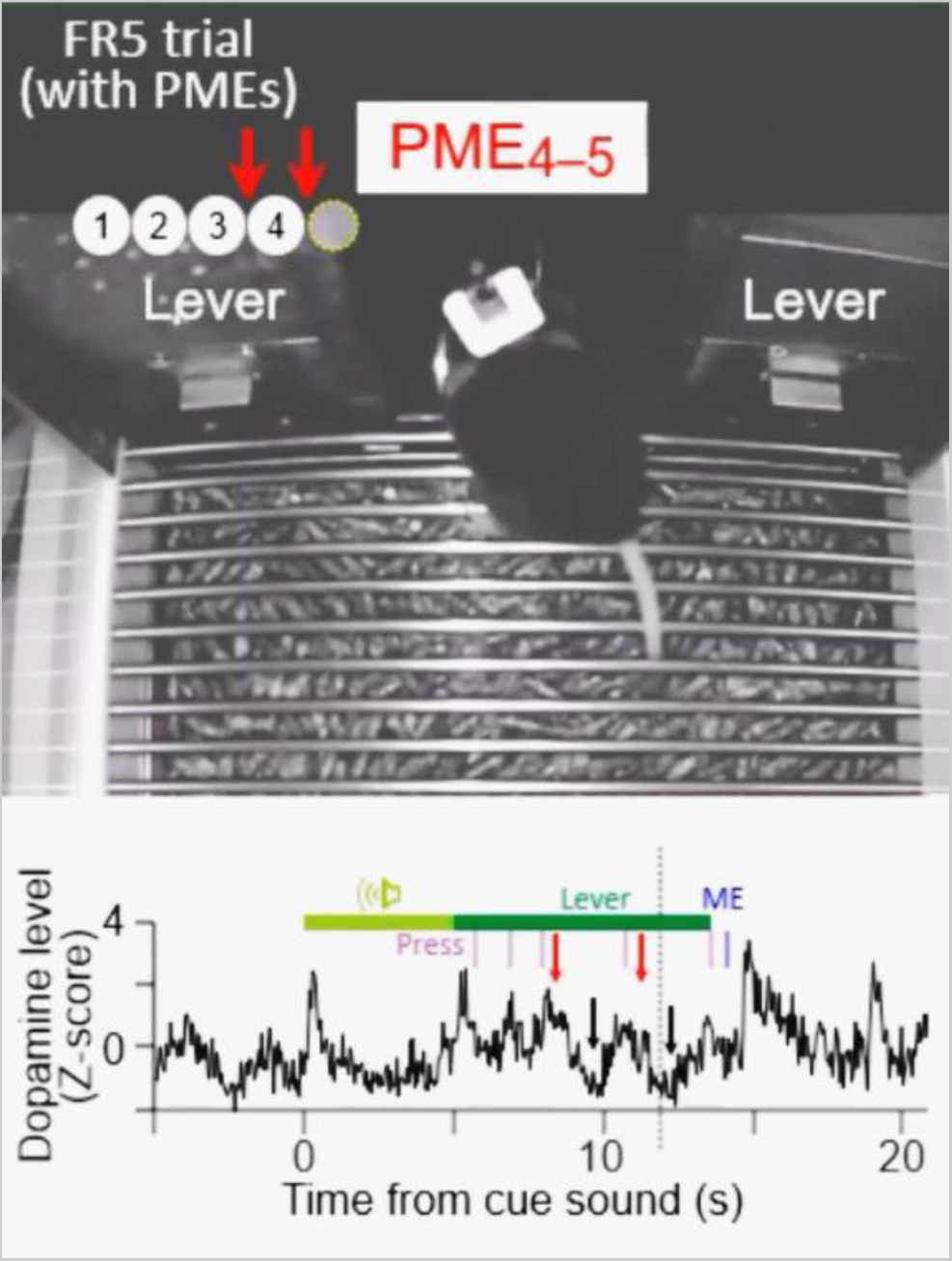
Phasic dopamine decrease upon a premature head entry into the magazine during the FR5 task. (Top) Representative behavior of a mouse engaging in the FR5 task. After the 5-s noise sound cue, reinforced (left) and non-reinforced (right) levers were presented. The mouse was rewarded at the magazine (white dashed line) after pressing the reinforced lever five times per trial. Of note, the mouse occasionally made a premature magazine entry (PME) between two consecutive lever presses. PME_4-5_, a PME between the fourth and the fifth lever presses. (Bottom) Simultaneous measurement of extracellular DA fluctuations. The vertical dotted line moves according to the camera frame timing. The DA level decreased transiently (black arrow just above the DA trace) following a PME (red arrow). We found that the amplitude of a DA dip upon a PME within a session increased with the increasing proximity to the task completion, e.g., a DA dip upon PME_4-5_ was, on average, larger than that upon PME_3-4_.

**Extended Data Figure 5-1.**
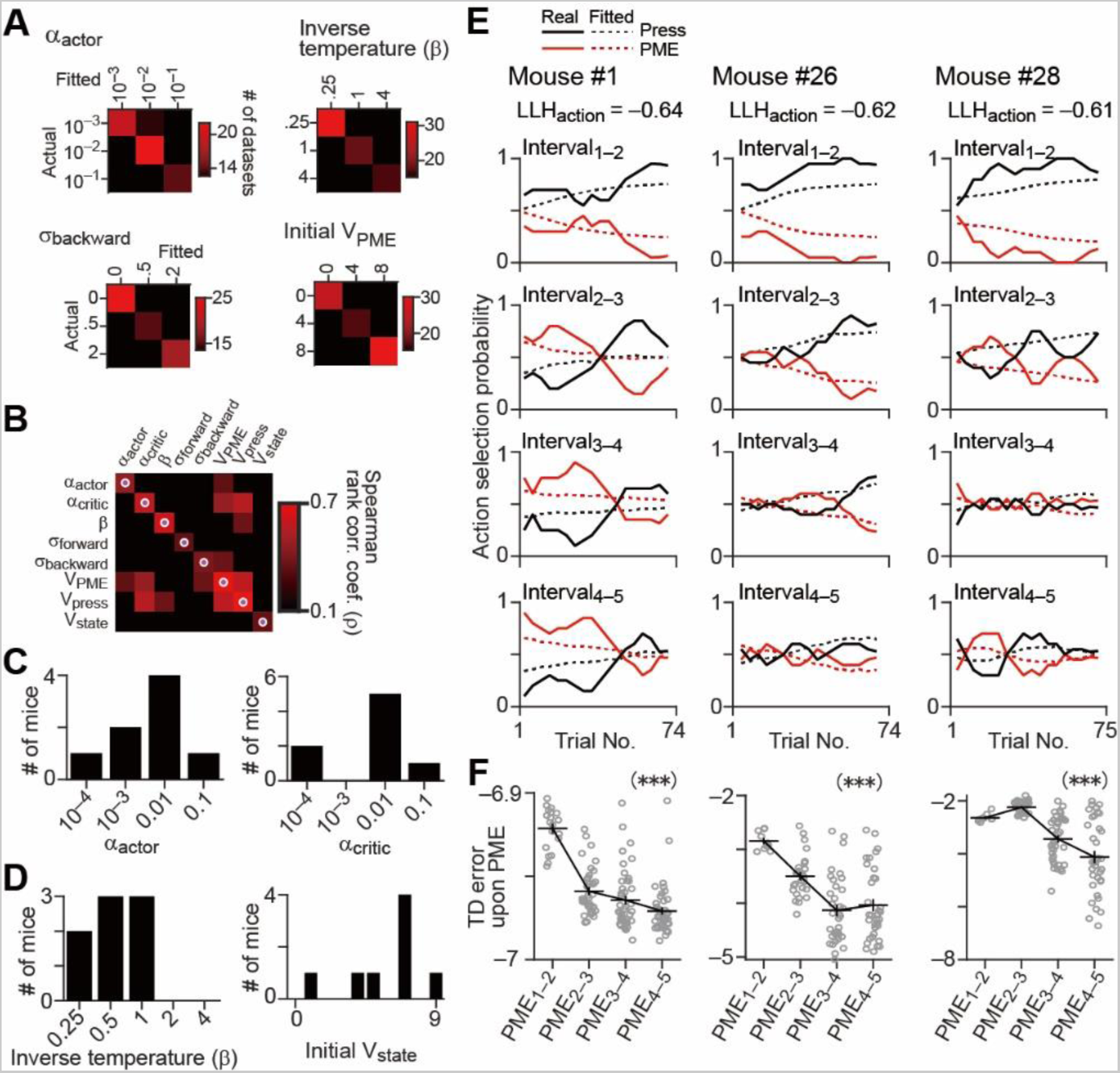
RL model simulation. (A, B) Parameter recovery and cross parameter correlations. A. Colormaps regarding individual parameters (α_actor_, inverse temperature (β), sigma for backward movement, and the initial action value of PME) showing the relationship between values with which synthesize data were made in the RL model and values recovered by fitting the model to those synthesized data. B. Colormap showing the Spearman’s rank correlation coefficients between simulated and recovered values with identical parameters (such as α_actor_ vs. α_actor_) or between recovered values for pairs of two different parameters (such as α_actor_ vs. initial V_PME_). Magenta dot: grid with the highest correlation coefficient in the corresponding row and column. (C) Group data showing the learning rates (α_actor_ [left] and α_critic_ [right]) used to calculate the TD errors for each step in a FR5 trial in the best-fitted model of the RL algorithm. (D) Related to ***Extended Data Fig. 5-1A*** but for β, the inverse temperature value used in the SoftMax function to compute the action selection probability (left), or for the initial state values assigned for the grids in the lever zone at the beginning of the FR5 schedule (right). (E) Related to Fig. 5D but showing the observed (solid lines) and simulated (dashed lines) action selection probability changes across the completed trials in the best-fitted model for other representative mice. (F) Same as Fig. 5H, showing statistical analyses in the TD error amplitude in the best-fitted model for other mice. The mice’s number corresponds to the above plots (***Extended Data Fig. 5-1E***). (Mouse 1, one-way ANOVA, *F*_(3,146)_ = 50.2, p = 3.1 × 10^−22^; Mouse 26, one-way ANOVA, *F*_(3,105)_ = 18.3, p = 1.4 × 10^−9^; Mouse 28, one-way ANOVA, *F*_(3,117)_ = 26.9, p = 3.2 × 10^−13^). Each open circle represents an individual event. Black line, mean ± SEM. *** p < 10^−8^.

## References

1. Amo R, Yamanaka A, Tanaka KF, Uchida N, Watabe-Uchida M (2020) A gradual backward shift of dopamine responses during associative learning. bioRxiv 2020.10.04.325324.

2. Bayer HM, Glimcher PW (2005) Midbrain Dopamine Neurons Encode a Quantitative Reward Prediction Error Signal. Neuron 47:129–141.

3. Bayer HM, Lau B, Glimcher PW (2007) Statistics of Midbrain Dopamine Neuron Spike Trains in the Awake Primate. J Neurophysiol 98:1428–1439.

4. Cachope R, Mateo Y, Mathur Brian N, Irving J, Wang H-L, Morales M, Lovinger David M, Cheer Joseph F (2012) Selective Activation of Cholinergic Interneurons Enhances Accumbal Phasic Dopamine Release: Setting the Tone for Reward Processing. Cell Reports 2:33–41.

5. Chéramy A, Romo R, Godeheu G, Baruch P, Glowinski J (1986) In vivo presynaptic control of dopamine release in the cat caudate nucleus—II. Facilitatory or inhibitory influence ofl-glutamate. Neuroscience 19:1081–1090.

6. Collins AGE, Frank MJ (2014) Opponent Actor Learning (OpAL): Modeling Interactive Effects of Striatal Dopamine on Reinforcement Learning and Choice Incentive. Psychol Rev 121:337–366.

7. Covey DP, Cheer JF (2019) Accumbal Dopamine Release Tracks the Expectation of Dopamine Neuron-Mediated Reinforcement. Cell Reports 27:481–490.e483.

8. Dabney W, Kurth-Nelson Z, Uchida N, Starkweather CK, Hassabis D, Munos R, Botvinick M (2020) A distributional code for value in dopamine-based reinforcement learning. Nature 577:671–675.

9. Dreyer JK, Herrik KF, Berg RW, Hounsgaard JD (2010) Influence of Phasic and Tonic Dopamine Release on Receptor Activation. J Neurosci 30:14273–14283.

10. Glowinski J, Chéramy A, Romo R, Barbeito L (1988) Presynaptic regulation of dopaminergic transmission in the striatum. Cell Mol Neurobiol 8:7–17.

11. Grieger JC, Choi VW, Samulski RJ (2006) Production and characterization of adeno-associated viral vectors. Nat Protoc 1:1412–1428.

12. Hamid AA, Pettibone JR, Mabrouk OS, Hetrick VL, Schmidt R, Weele CM, Kennedy RT, Aragona BJ, Berke JD (2016) Mesolimbic dopamine signals the value of work. Nat Neurosci 19:117–126.

13. Hart AS, Rutledge RB, Glimcher PW, Phillips PEM (2014) Phasic Dopamine Release in the Rat Nucleus Accumbens Symmetrically Encodes a Reward Prediction Error Term. J Neurosci 34:698–704.

14. Heien M, Khan AS, Ariansen JL, Cheer JF, Phillips PEM, Wassum KM, Wightman MR (2005) Real-time measurement of dopamine fluctuations after cocaine in the brain of behaving rats. Proc Natl Acad Sci USA 102:10023–10028.

15. Howe MW, Dombeck DA (2016) Rapid signalling in distinct dopaminergic axons during locomotion and reward. Nature 535:505–510.

16. Iino Y, Sawada T, Yamaguchi K, Tajiri M, Ishii S, Kasai H, Yagishita S (2020) Dopamine D2 receptors in discrimination learning and spine enlargement. Nature 579:555–560.

17. Kahneman D, Tversky A (1979) On the interpretation of intuitive probability: A reply to Jonathan Cohen. Cognition 7:409–411.

18. Ko D, Wanat MJ (2016) Phasic Dopamine Transmission Reflects Initiation Vigor and Exerted Effort in an Action- and Region-Specific Manner. J Neurosci 36:2202–2211.

19. Mathiesen SN, Lock JL, Schoderboeck L, Abraham WC, Hughes SM (2020) CNS Transduction Benefits of AAV-PHP.eB over AAV9 Are Dependent on Administration Route and Mouse Strain. Molecular Therapy - Methods & Clinical Development 19:447–458.

20. Mohebi A, Pettibone JR, Hamid AA, Wong J-MT, Vinson LT, Patriarchi T, Tian L, Kennedy RT, Berke JD (2019) Dissociable dopamine dynamics for learning and motivation. Nature 570:65–70.

21. Moutoussis M, Bentall RP, Williams J, Dayan P (2009) A temporal difference account of avoidance learning. Network: Comput Neural Syst 19:137–160.

22. Nakahara H, Itoh H, Kawagoe R, Takikawa Y, Hikosaka O (2004) Dopamine Neurons Can Represent Context-Dependent Prediction Error. Neuron 41:269–280.

23. Natsubori A, Tsutsui-Kimura I, Nishida H, Bouchekioua Y, Sekiya H, Uchigashima M, Watanabe M, d’Exaerde Ad, Mimura M, Takata N, Tanaka KF (2017) Ventrolateral Striatal Medium Spiny Neurons Positively Regulate Food-Incentive, Goal-Directed Behavior Independently of D1 and D2 Selectivity. J Neurosci 37:2723–2733.

24. Neumann Jv, Morgenstern O (1945) Theory of Games and Economic Behavior. J Philos 42:550–554.

25. Niv Y, Daw ND, Joel D, Dayan P (2007) Tonic dopamine: opportunity costs and the control of response vigor. Psychopharmacology (Berl) 191:507–520.

26. O’Doherty JP, Dayan P, Friston K, Critchley H, Dolan RJ (2003) Temporal Difference Models and Reward-Related Learning in the Human Brain. Neuron 38:329–337.

27. Phillips PEM, Stuber GD, Heien MLAV, Wightman RM, Carelli RM (2003) Subsecond dopamine release promotes cocaine seeking. Nature 422:614–618.

28. Robinson DL, Venton BJ, Heien MLAV, Wightman RM (2003) Detecting Subsecond Dopamine Release with Fast-Scan Cyclic Voltammetry in Vivo. Clin Chem 49:1763–1773.

29. Rodeberg NT, Sandberg SG, Johnson JA, Phillips PEM, Wightman MR (2017) Hitchhiker’s Guide to Voltammetry: Acute and Chronic Electrodes for in Vivo Fast-Scan Cyclic Voltammetry. ACS Chem Neurosci 8:221–234.

30. Schultz W, Dayan P, Montague RP (1997) A Neural Substrate of Prediction and Reward. Science 275:1593–1599.

31. Starkweather C, Gershman SJ, Uchida N (2018) The Medial Prefrontal Cortex Shapes Dopamine Reward Prediction Errors under State Uncertainty. Neuron 98:616–629.e616.

32. Starkweather CK, Uchida N (2021) Dopamine signals as temporal difference errors: recent advances. Curr Opin Neurobiol 67:95–105.

33. Sun F, Zeng J, Jing M, Zhou J, Feng J, Owen SF, Luo Y, Li F, Wang H, Yamaguchi T, Yong Z, Gao Y, Peng W, Wang L, Zhang S, Du J, Lin D, Xu M, Kreitzer AC, Cui G, Li Y (2018) A Genetically Encoded Fluorescent Sensor Enables Rapid and Specific Detection of Dopamine in Flies, Fish, and Mice. Cell 174:481–496.e419.

34. Sun F, Zhou J, Dai B, Qian T, Zeng J, Li X, Zhuo Y, Zhang Y, Tan K, Feng J, Dong H, Qian C, Lin D, Cui G, Li Y (2020) New and improved GRAB fluorescent sensors for monitoring dopaminergic activity in vivo. bioRxiv 2020.03.28.013722.

35. Suri RE, Schultz W (2001) Temporal Difference Model Reproduces Anticipatory Neural Activity. Neural Comput 13:841–862.

36. Threlfell S, Lalic T, Platt Nicola J, Jennings Katie A, Deisseroth K, Cragg Stephanie J (2012) Striatal Dopamine Release Is Triggered by Synchronized Activity in Cholinergic Interneurons. Neuron 75:58–64.

37. Tian J, Uchida N (2015) Habenula Lesions Reveal that Multiple Mechanisms Underlie Dopamine Prediction Errors. Neuron 87:1304–1316.

38. Tobler PN, Fiorillo CD, Schultz W (2005) Adaptive Coding of Reward Value by Dopamine Neurons. Science 307:1642–1645.

39. Tsutsui-Kimura I, Natsubori A, Mori M, Kobayashi K, Drew MR, d’Exaerde AdK, Mimura M, Tanaka KF (2017a) Distinct Roles of Ventromedial versus Ventrolateral Striatal Medium Spiny Neurons in Reward-Oriented Behavior. Curr Biol 27:3042–3048.e3044.

40. Tsutsui-Kimura I, Takiue H, Yoshida K, Xu M, Yano R, Ohta H, Nishida H, Bouchekioua Y, Okano H, Uchigashima M, Watanabe M, Takata N, Drew MR, Sano H, Mimura M, Tanaka KF (2017b) Dysfunction of ventrolateral striatal dopamine receptor type 2-expressing medium spiny neurons impairs instrumental motivation. Nat Commun 8:14304.

41. Yoshida K, Drew MR, Mimura M, Tanaka KF (2019) Serotonin-mediated inhibition of ventral hippocampus is required for sustained goal-directed behavior. Nat Neurosci 22:770–777.

42. Yoshida K, Tsutsui-Kimura I, Kono A, Yamanaka A, Kobayashi K, Watanabe M, Mimura M, Tanaka KF (2020) Opposing Ventral Striatal Medium Spiny Neuron Activities Shaped by Striatal Parvalbumin-Expressing Interneurons during Goal-Directed Behaviors. Cell Reports 31:107829.

43. Zhou F-M, Liang Y, Dani JA (2001) Endogenous nicotinic cholinergic activity regulates dopamine release in the striatum. Nat Neurosci 4:1224–1229.

